# Ancient origin of the rod bipolar cell pathway in the vertebrate retina

**DOI:** 10.1101/2023.09.12.557433

**Authors:** Ayana M Hellevik, Philip Mardoum, Joshua Hahn, Yvonne Kölsch, Florence D D’Orazi, Sachihiro C. Suzuki, Leanne Godinho, Owen Lawrence, Fred Rieke, Karthik Shekhar, Joshua R Sanes, Herwig Baier, Tom Baden, Rachel O Wong, Takeshi Yoshimatsu

**Affiliations:** Department of Biological Structure, University of Washington, Seattle, WA 98195, USA; Department of Chemical and Biomolecular Engineering; Helen Wills Neuroscience Institute; Vision Sciences Graduate Program; California Institute of Quantitative Biosciences (QB3), University of California Berkley, Berkeley, CA 94720, USA; Department of Molecular & Cellular Biology and Center for Brain Science, Harvard University, Cambridge, MA 02138, USA; Max Planck Institute for Biological Intelligence, Department Genes – Circuits – Behavior, 82152 Martinsried, Germany; Institute of Neuronal Cell Biology, Technische Universität München, 80802 Munich, Germany; Department of Physiology and Biophysics, University of Washington, Seattle, WA 98195, USA; Vision Science Center, University of Washington, Seattle, WA 98195, USA; Biological Systems and Engineering Division, Lawrence Berkeley National Laboratory, Berkeley, CA 94720, USA; School of Life Sciences, University of Sussex, Brighton, BN1 9QG, UK; Institute of Ophthalmic Research, University of Tübingen, Tübingen, 72076, Germany; Department of Ophthalmology & Visual Sciences, Washington University in St Louis School of Medicine, St Louis, MO 63110, USA; BioRTC, Yobe State University, Damatsuru, Yobe 620101, Nigeria

**Author notes:** These authors contributed equally to this study (Ayana M Hellevik, Philip Mardoum, Joshua Hahn). Lead contact: Takeshi Yoshimatsu.

**Keywords:** Rod, Rod bipolar cells, Retina, Evolution

## Abstract

Vertebrates rely on rod photoreceptors for vision in low-light conditions^1^. Mammals have a specialized downstream circuit for rod signaling called the primary rod pathway, which comprises specific cell types and wiring patterns that are thought to be unique to this lineage^2–6^. Thus, it has been long assumed that the primary rod pathway evolved in mammals^3, 5–7^. Here, we challenge this view by demonstrating that the mammalian primary rod pathway is conserved in zebrafish, which diverged from extant mammals ∼400 million years ago. Using single-cell RNA-sequencing, we identified two bipolar cell (BC) types in zebrafish that are related to mammalian rod BCs (RBCs) of the primary rod pathway. By combining electrophysiology, histology, and ultrastructural reconstruction of the zebrafish RBCs, we found that, like mammalian RBCs^8^, both zebrafish RBC types connect with all rods and red-cones in their dendritic territory, and provide output largely onto amacrine cells. The wiring pattern of the amacrine cells post-synaptic to one RBC type is strikingly similar to that of mammalian RBCs. This suggests that the cell types and circuit design of the primary rod pathway may have emerged before the divergence of teleost fish and amniotes (mammals, bird, reptiles). The second RBC type in zebrafish, which forms separate pathways from the first RBC type, is either lost in mammals or emerged in fish to serve yet unknown roles.

**Highlights:**

- Zebrafish have two rod bipolar cell types (RBC1/2).
- Synaptic connectivity of RBC1 resembles that of the mammalian RBCs.
- The primary rod pathway therefore probably evolved more than 400 million years ago.
- The second zebrafish RBC type, RBC2, forms a separate pathway from RBC1.

## INTRODUCTION

Rod photoreceptors of the vertebrate retina are capable of detecting very dim light, down to individual photons^9–14^. The mammalian cell types and circuitry that convey rod-driven signals, called the primary rod pathway, were identified and defined in the cat retina about 50 years ago^15^. Since its initial characterization, this rod pathway has been examined extensively in many mammalian species including in humans, where it consistently uses homologous cell types and connectivity patterns: Rod bipolar cells (RBCs), as well as A2 and A17 amacrine cells^3, 5, 6, 16, 17^. The RBC is a molecularly, structurally and functionally distinct retinal bipolar cell type that receives input from all rods within its dendritic field and is predominantly driven by rods^8, 18–21^. In mammals, all other bipolar cell types receive most of their photoreceptor input from cones, which operate in daylight conditions. In contrast, the bipolar cells that have been characterized in non-mammals lack clear distinctions regarding the ratio of rod and cone inputs^14, 22^. Thus, it is not surprising that the RBC is thought to be unique to mammals, a notion that has led to the prevailing view that the rod pathway evolved separately in this class^3, 5–7^. However, the molecular, structural and functional signatures that together define bipolar cell types in mammals are largely unknown in non-mammals, so it remains unclear whether the signatures characteristic of the mammalian RBC and its downstream pathway are present in non-mammals. To address this issue, we focused on zebrafish, a species that allows transgenic labeling of neuronal populations, to analyze single-cell transcriptomics, histology, physiology and circuit reconstructions of genetically-defined retinal bipolar cells.

There are more than a dozen BC types in mammals that are diverse in morphology, connectivity and molecular profiles^23–27^, but can be classified into two main groups: ON BCs that depolarize and OFF BCs that hyperpolarize in response to increases in luminance^28^. RBCs are one type of ON BCs that are distinct in many ways from all other BCs, which mainly connect with cones and are called cone BCs (CBCs) here. Transcriptionally, mammalian RBCs can be distinguished from CBCs by the expression of protein kinase C-alpha (PKCα)^29^. Morphologically, the axon terminals of RBCs are generally larger than those of CBCs and end in the innermost layer of the inner plexiform layer (IPL). The synaptic arrangement of the RBC axons differs from the common synaptic arrangement of most cone bipolar cells. Whereas CBC axons directly synapse onto the retinal output neurons, retinal ganglion cells (RGCs), along with a plethora of amacrine cells (ACs), RBCs predominately form a ’dyadic’ synapse with two types of inhibitory amacrine cells, small field A2 (or A-II) and large-field A17 ACs^15, 18, 19, 30^. The A17 AC almost exclusively makes reciprocal feedback synapses onto RBC axon terminals^31^. In contrast, A2 ACs receive numerous synapses from RBCs (∼40 synapses per RBC in mice), but do not provide feedback onto the RBCs^18, 32^. These RBC to A2 AC synapses are the critical sites for the amplification and gain control of rod signals^33–35^. Rod signals are eventually relayed to RGCs by connections from A2 ACs on CBCs, which split rod signals into ON and OFF channels via sign-conserving gap junctions with ON CBCs and inhibitory synapses with OFF CBCs^15, 36, 37^.

Here, by analyzing single-cell transcriptomic profiles of zebrafish BCs, we discovered two BC types, RBC1 and RBC2, with molecular signatures similar to those of mammalian RBCs. Using transgenic zebrafish lines that express a fluorescent protein in RBC1 or RBC2 cells, we identified the inputs and outputs of RBC1 and RBC2. We found that both zebrafish RBC types connect with all rods and red cones (or longwave-length sensitive, LWS, cones) inside their dendritic fields. We further reconstructed the downstream circuits of both BC types using serial block-face electron microscopy and found that RBC1 predominantly synapses onto three morphological types of ACs. The circuit diagrams and synaptic arrangements of two of the ACs closely resemble those of the mammalian A2 and A17 ACs. In contrast, RBC2 mainly connects to a different set of ACs, which does not include A2-like ACs. These results suggest that (i) zebrafish possess two separate pathways for processing rod signals, and that (ii) one of these is similar to the rod –> RBC –> A2 AC –> CBC –> RGC pathway found in mammals. We conclude that the primary rod pathway emerged >400 million years ago, before the divergence of teleosts and mammals in the Devonian.

## RESULTS

### Two zebrafish BC types are transcriptionally analogous to the mammalian RBCs

We first determined the transcriptional similarity between each zebrafish BC type and the mammalian RBCs by using single-cell RNA-sequencing (scRNA-seq) in adult zebrafish. BCs were isolated using a fluorescent marker in the *Tg(vsx1:GFP)^nns^*^5^ transgenic line, in which all BCs express GFP^38^ (Fig. 1a). Clustering analysis of 19492 high-quality single cell transcriptomes identified 23 molecularly distinct BC clusters (Fig. 1b,c). To identify the clusters most similar to mammalian RBCs, we performed a hierarchical clustering analysis based on average transcriptomic profiles (Fig. 1d) and combined this with the expression patterns of marker genes identified in mice to tentatively annotate each cluster as ON CBC, OFF CBC or RBC (Fig. 1e; ^27^). In mice, RBCs are clearly separated from CBCs at the first dendrogram bifurcation^27^. Similarly, the first dendrogram bifurcation separates two BCs from the other BCs in zebrafish. In contrast to mice, however, the zebrafish RBC clade contained two molecularly distinct clusters 14 (c14) and 19 (c19) (Fig. 1d). We observed that *prkca* (the gene encoding PKCα), a common marker of mammalian RBCs, is only highly expressed in c14. However, both c14 and c19 specifically express *gramd1b*, which is an RBC-specific marker in mice. In addition to these genetic signatures similar to mammalian RBCs, both c14 and c19 clusters express neurotransmitter receptor, *grm6a* and *grm6b,* and it’s downstream signaling molecules, *trpm1a, trpm1b*, *nyx* and *rgs11,* which are essential for mediating rod inputs in mammals^39^ (Fig. S1). Therefore, we hypothesized that zebrafish, unlike mice, may possess two RBC types, which we call RBC1 (c14) and RBC2 (c19).

**Figure 1.**
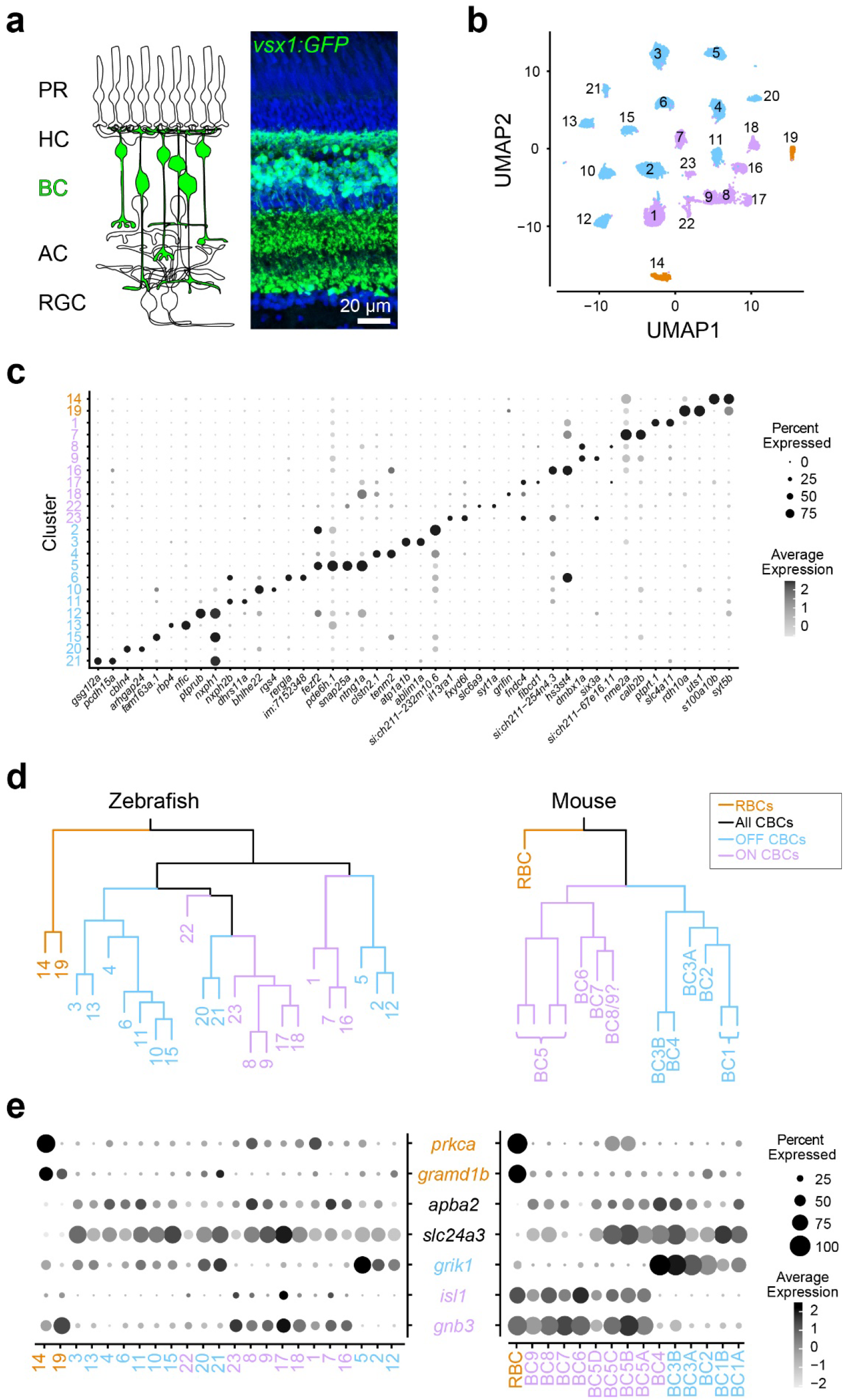
Comparison of single-cell gene expressions identified two possible rod bipolar cells in zebrafish. **a**, Schematic representation of retinal circuits (left) and an image of a retinal slice from Tg(vsx1:GFP)^nns5^ transgenic adult zebrafish (right). GFP expression in all bipolar cells (BCs). Nuclei was stained by DAPI. PR: photoreceptor, HC: horizontal cell, AC: amacrine cell, RGC: retinal ganglion cell. **b**, 2D visualization of single-cell clusters using Uniform Manifold Approximation (UMAP)^40^. Individual points correspond to single cells colored according to cluster identity. **c**, Marker genes for each cluster. **d**, Agglomerative hierarchical clustering of average gene signatures of clusters using the correlation metric and complete linkage. BC subclasses (colors) were assigned based on the known marker expressions shown in **e**. **e**, Gene expression patterns of known BC subclass markers in BC clusters. The size of each circle depicts the percentage of cells in the cluster in which the marker was detected (≥1 UMI), and its contrast depicts the scaled average expression level of cells within the cluster in **c,e**. Data for mouse is from Shekhar K, et al., 2016, Cell.

### RBC1 and RBC2 morphologies resemble mammalian RBCs

We next determined the morphological similarities of RBC1 and RBC2 with mammalian RBCs. In mammals, RBC axons arborize in the innermost layer of the IPL^40^. By screening our zebrafish transgenic lines, we identified two lines, *Tg(vsx1:memCerulean)^q19^* (*vsx1:memCer)* and *Tg(vsx2:memCerulean)^wst01^* (*vsx2:memCer)*, that each label BCs with axon terminals in the innermost layer of the IPL (Fig. 2). Fluorescent *in situ* hybridization for the identified gene markers, *s100a10b* and *uts1,* which are selectively expressed by RBC1 and RBC2 (Fig. 1c), revealed that *vsx1:memCer* and *vsx2:memCer* label RBC1 and RBC2, respectively (Fig. 2a,b). We also observed that dendritic arbors of both RBC1 and RBC2 cover the retina in a non-overlapping manner, an arrangement called ’tiling’ that is considered a hallmark of a BC type (Fig. 2e,f)^41^. Therefore, both RBC1 and RBC2 represent single bipolar types that transcriptionally and morphologically resemble mammalian RBCs.

**Figure 2.**
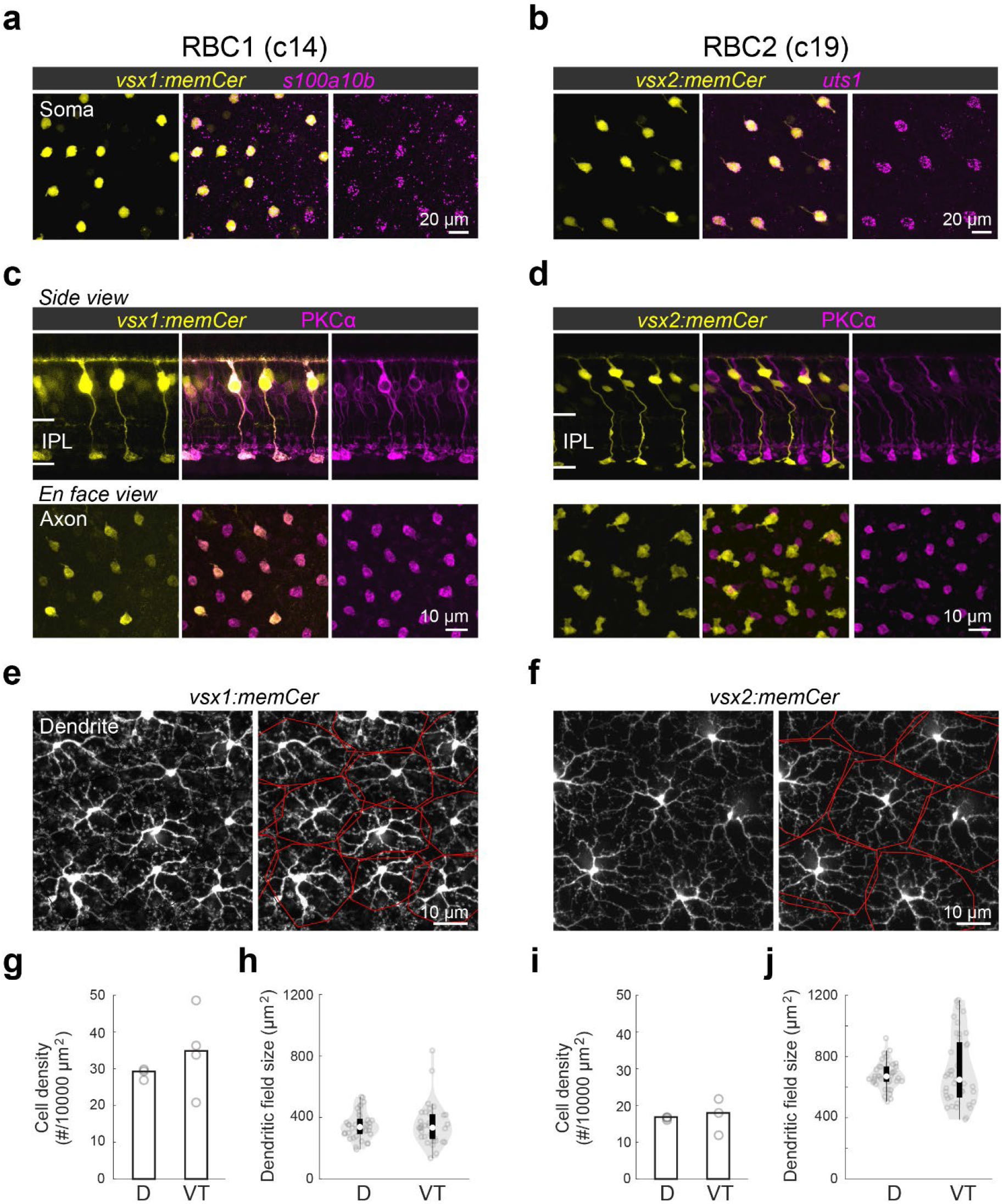
Transgenic labeling of cluster 14 and 19 revealed morphological features of these BCs. **a,b**, En face view of retinal flat mount at the inner nucleus layer level. Cerulean fluorescent expression (colored yellow) transgenic lines, Tg(vsx1:memCerulean)^q19^ (vsx1:memCer) in **a** and Tg(vsx2:memCerulean)^wst01^ (vsx2:memCer) in **b**. vsx1:memCer and vsx2:memCer BCs are positive for cluster specific genes, s100a10b and uts1, respectively, which are detected using in situ hybridization chain reaction ^43^. **c,d**, Side views of the labeled cells and the distribution patterns of their axon terminals in en face views of retinal flat mounts for RBC1 (**c**) and RBC2 (**d**) BCs. Immunolabeling for PKCα is in magenta. IPL: inner plexiform layer. Note that not all PKC-immunoreactive cells are apparent in this image of the vsx1:memCer line, due to the incomplete labeling of this line. **e,f**, Dendritic tiling of RBC1 (**e**) and RBC2 (**f**) in en face view of retinal flat mounts at the outer plexiform layer level. Dendritic territories are marked by the red boundaries. **g-j**, Mean cell densities of RBC1 (**g**, n=3 and 4 for D and VT, respectively) and RBC2 (**I**, n=3 for both D and VT) BCs in different regions of the retina. Box and violin plots of dendritic field sizes of RBC1 (h, n=37 and 33 or D and VT, respectively) and RBC2 (j, n=40 and 41 or D and VT, respectively) BCs. White filled circles are medians. Grey circles indicate individual cells. D: dorsal, VT: ventrotemporal.

We observed slight variations in morphology and molecular expression between RBC1 and RBC2. The axon terminal of RBC1 is relatively spherical, similar to mammalian RBCs, in contrast to the ‘flat-footed’ axonal ending of RBC2 (Fig. 2c,d). RBC1 were immunoreactive for PKCα (Fig. 2c), whereas RBC2 were not (Fig. 2d), consistent with the difference in their *prkca* expression (Fig. 1d). In addition, their abundance differed: RBC1s were more densely packed than RBC2s (*p=0.0052*, Mann-Whitney two-tailed U test) (Fig. 2g,i). This difference in the densities is unlikely due to regional variations as both RBCs are present in the dorsal and ventral-temporal retina at similar densities (Fig. 2g,i and Fig. S2). The dendritic field sizes of the two RBC types were inversely related to their cell density, consistent with their tiling arrangement (Fig. 2g-j).

### Both RBC1 and RBC2 connect with all rods and red-cones in their dendritic territory

If RBC1 and RBC2 are authentic RBCs, they should synapse preferentially with rods. Using 4C12 antibodies to label rods, we found that the majority of the dendritic tips of both RBC types (RBC1: 84±3.9%, RBC2: 78±2.9%) contacted with rod spherules (Fig. 3a-d). We also found that some dendritic tips were not associated with rods (Fig. 3a-d). Using transgenic lines to label specific cone types, we identified that dendritic tips of both BC types contacted red-cones (or long-wavelength sensitive cones), labeled in the *Tg(trb2:tdtomato)^q22^* line (Fig. 3a-e,g). Furthermore, both types connected with nearly all rods and red cones within their dendritic fields (Fig. 3f,h). We did not observe any dendritic tips that were not associated with either rods or red cones (Fig. 3e,g), indicating that they receive few if any inputs from the other cone types, which include green, blue, and violet cones^42, 43^. Therefore, both RBC1 and 2 receive predominant rod input and share specificity for red cones among cones (Fig. 3e,g).

**Figure 3.**
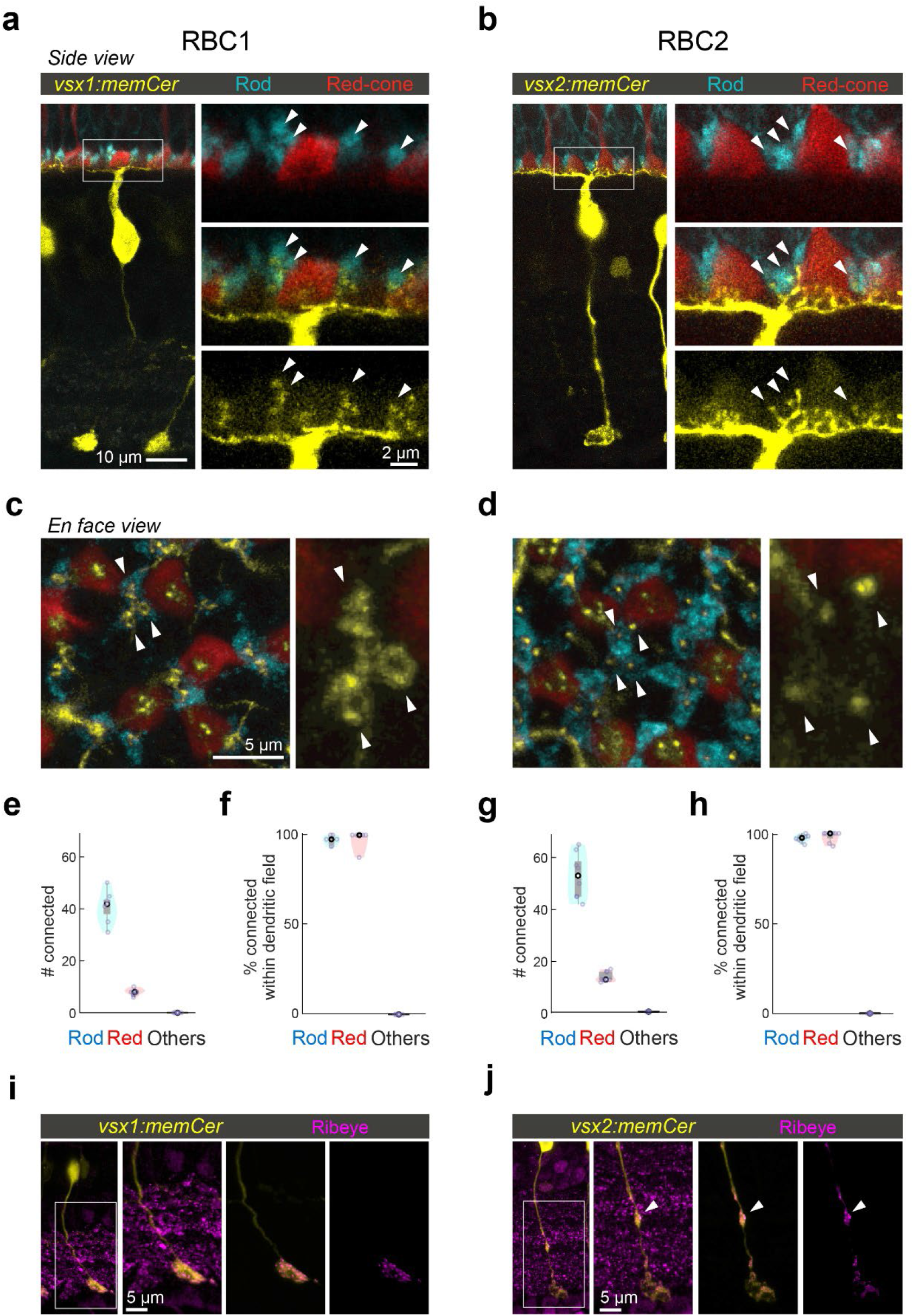
RBC1 and RBC2 connect to rods and red cones but differ in dendritic and axonal synaptic arrangements. **a,b**, Dendritic tips invaginating the rod and red cone axon terminals, visualized in retinal slices from (a) Tg(vsx1:memCerulean:trb2:tdtomato)^q19,q22^ and (b) Tg(vsx2:memCerulean:trb2:tdtomato)^wst01,q22^ adult zebrafish. Rods were immunolabeled using 4C12 antibody. **c,d**, Doughnut and simple dendritic tip structures at rod terminals (arrow heads) in RBC1 and RBC2, respectively. **e-h**, Box and violin plots of RBC1 (**e,f**, n=7) and RBC2 (**g,h**, n=8) connectivity with photoreceptors. **i,j**, Distribution of ribbon synapses in the RBC1 (**i**) and 2 (**j**) axons. Ribbons were immunolabeled by anti-ribeye antibody. Ribeye signals outside the axons were digitally masked out in the right two images. RBC2 axon harbors a ribbon containing distal bouton in the OFF layer (arrow head).

Interestingly, the dendritic tips of RBC1 and RBC2 terminating in rod spherules differed in structure (Fig. 3c,d). Specifically, the dendrites of RBC1 invaginating rod spherules appeared to form a horseshoe or ‘doughnut’ ending, whereas those of RBC2 ended in a simpler arrangement (Fig. 3c,d). The larger surface area of RBC1’s dendritic tips at rod terminals may increase the sensitivity to rod inputs in this BC type compared to RBC2. We also observed differences in the distal axonal boutons of RBC1 and RBC2 (Fig. 3i,j). While axons of both types terminate close to the ganglion cells in the IPL, RBC2 axons have a bouton in the OFF layer of the IPL, next to the boundary with the ON layer (Fig. 3j). These distal boutons are likely pre-synaptic sites as they contain the pre-synaptic protein, Ribeye (Fig. 3j). These differences in the dendritic tip and axon bouton shapes between RBC1 and RBC2 suggest that, while both BCs receive input from the same combination of photoreceptor types, they may serve distinct visual functions.

### RBC1 receives rod inputs via mGluR6 receptors

We next asked whether zebrafish RBCs receive functional rod input via the metabotropic glutamate receptor mGluR6 as seen in mammalian RBCs. We first investigated the expression of mGluR6 in RBC1 and RBC2 dendritic tips at rod spherules. Super-resolution imaging of mGluR6 immunolabeling in *vsx1:memCerulean and vsx2:memCerulean* retinas showed that the dendritic tips of RBC1, but not RBC2, robustly overlapped with mGluR6 immunoreactivity at contacts with rod spherules (Fig. 4a,b). These findings are consistent with the transcriptional profiles, which showed that RBC1 expresses higher mRNA levels of *grm6a* and *grm6b*, which encodes mGluR6, than RBC2 (Fig. S1).

**Figure 4.**
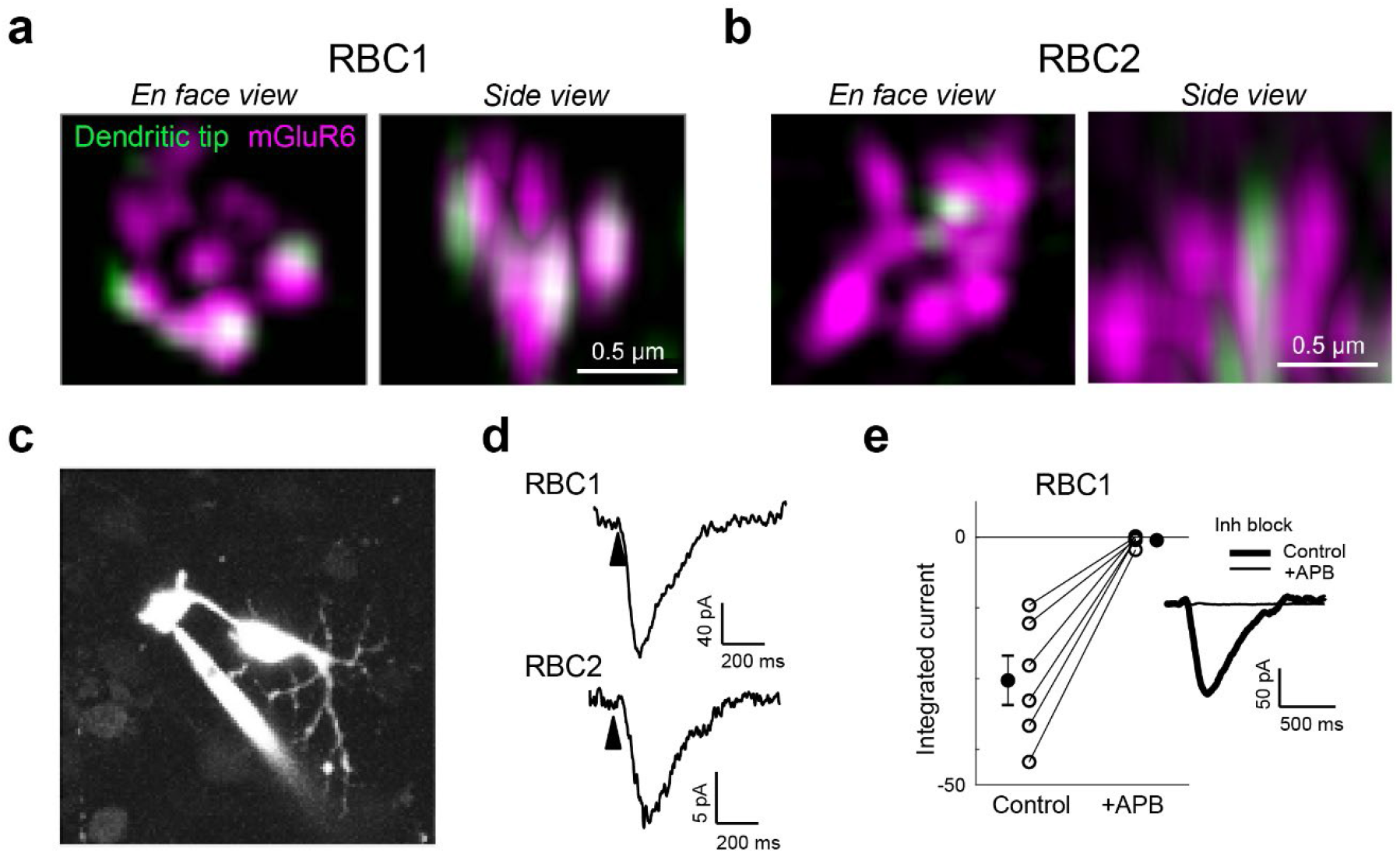
Rod input to RBC1 is mediated by mGluR6 receptors. **a,b**, Colocalization of mGluR6 and RBC1 (**a**) and RBC2 (**b**) dendritic tips at rod terminals visualized by structured illumination microcopy. **c**, Whole-cell patch clamping of a RBC1 axon terminal, visualized by dye-filling Alexa Fluor 594). **d**, Voltage responses of RBC1 and RBC2 after a cone activating light flash (arrow heads). **e,** Population data of RBC1 responses to rod activating light flashes with and without the group III metabotropic glutamate receptor agonist, APB (6-(2-aminopropyl)benzofuran). Filled circles: mean; error bars, S.D.; open circles, individual cells. Traces on the right are an example of the cell’s light evoked response before and during APB bath application. Inhibitory neurotransmitter receptors were blocked (inh lock) by a bath application of gabazine, strychnine, and TPMPA ((1,2,5,6-Tetrahydropyridin-4-yl)methylphosphinic acid).

We then used electrophysiological recordings to ask whether mGluR6 mediates rod input to the zebrafish RBCs. We prepared retinal wholemounts that preserve synaptic connections in the outer retina, and performed whole-cell patch-clamp recordings on the axon terminals of RBC1 and RBC2 (Fig. 4c). Both RBC1 (n = 10) and RBC2 cells (n = 3) exhibited ON responses to a cone-activating flash (red LED), confirming the successful patch-clamp recordings of light responses in these BCs and demonstrating that both cell types are ON cells (Fig. 4d), consistent with the position of their axonal arbors (Fig. 2c,d).

Although measuring rod-mediated responses from RBC2 was infeasible for technical reasons (see Methods), we were successful in recording rod responses from RBC1. We were therefore able to ask whether these responses mediated by mGluR6. We presented rod-isolating dim blue flashes (10 ms) before and after introducing the mGluR6 receptor agonist 6-(2-aminopropyl)benzofuran (APB) to the perfusion solution. To isolate excitatory inputs to the cell, all recordings were performed near the reversal potential for chloride-mediated conductances (∼-60 mV) and in the presence of inhibitory receptor blockers, gabazine, strychnine, and TPMPA ((1,2,5,6-Tetrahydropyridin-4-yl)methylphosphinic acid). Our results showed that nearly all rod inputs were blocked in the presence of APB, indicating that, like mammalian RBCs, mGluR6 mediates rod input to RBC1 (Fig. 4e).

### Both RBCs primarily synapse onto amacrine cells

To determine the synaptic targets of RBC1 and RBC2, we reconstructed their connectomes using serial block face scanning electron microscopy (SBFSEM). In the reconstructions, we observed an array of large BC axon terminals in the innermost layer of the IPL, which are characteristics of RBC1 and RBC2 axons (Fig. S3a,b). To confirm that these large axon terminals belong to RBC1 and RBC2, we reconstructed dendrites of some of these BCs (Fig. 5a). Consistent with our observations in light microscopic experiments (Fig. 3a-d), the large axon BCs predominantly connect with rods.

**Figure 5.**
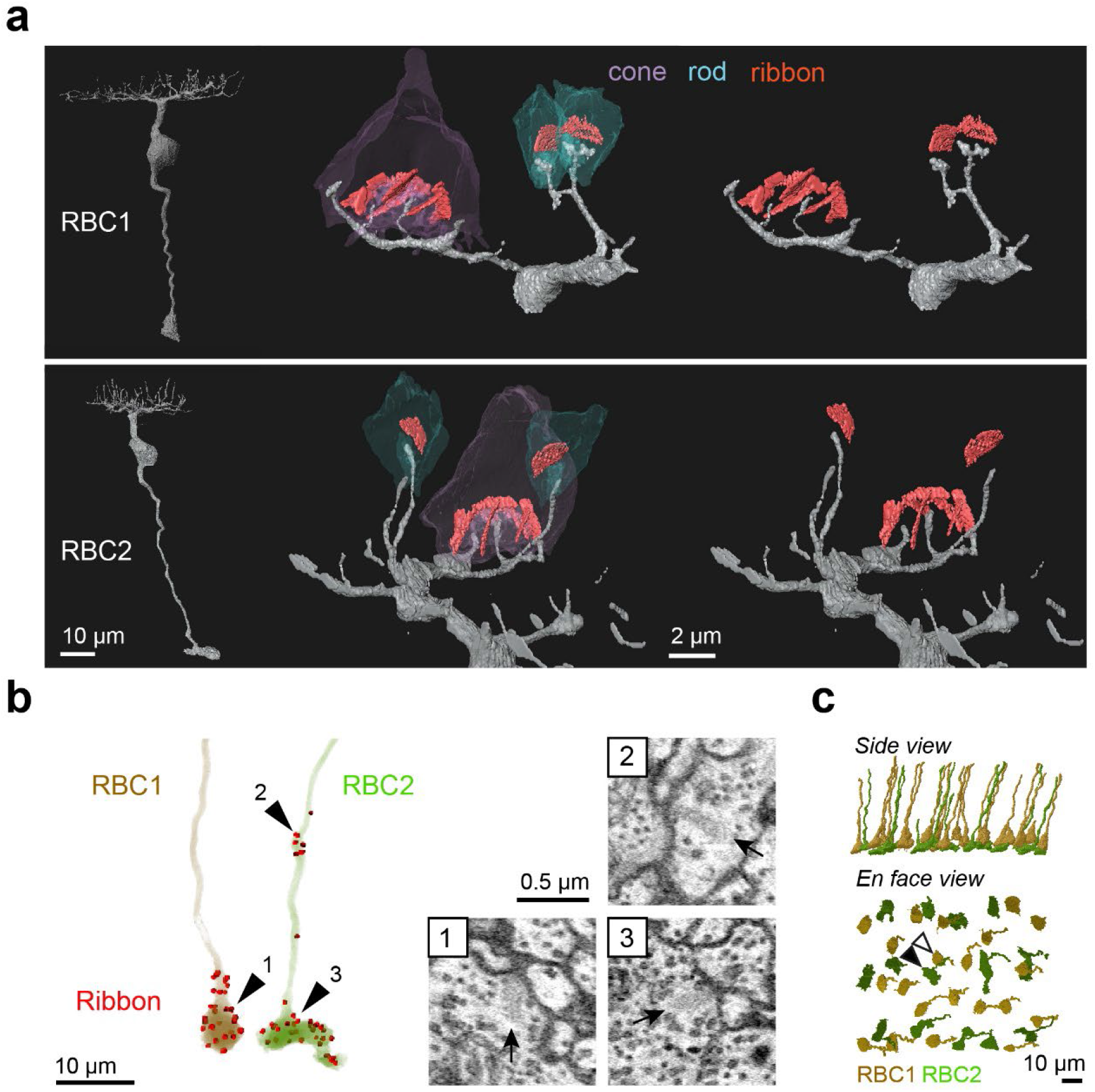
Identification of RBC1 and RBC2 in a SBFSEM volume. **a**, Reconstructions of a RBC1 and a RBC2, and zoomed-in images of their dendritic tips at rod and cone terminals. Ribbons in the rod and cones are painted red. **b**, Ribbon synapse distributions in a RBC1 and a RBC2. The locations of ribbon synapses are marked in red. Arrow heads indicate the locations of example ribbon synapses (arrows) shown in the insets. **c**, Reconstruction of all RBC1s and RBC2s in the EM volume. Postsynaptic neurons of a centrally located RBC1 (open arrow head) and RBC2 (closed arrow head) were reconstructed in Fig. 6, 7, and S5-9.

We then reconstructed all the large axons in the SBFSEM image volume. To distinguish RBC2 from RBC1, we used the ribbon containing axonal distal bouton in the OFF layer as a proxy for RBC2 (Fig. 3i,j and Fig. 5b). These reconstructions revealed the regular mosaic arrangements of both presumed RBC1 and RBC2 (Fig. 5c), indicating that we identified most, if not all, presumed RBC1 and RBC2 in the EM volume. Using this criterion, we also verified that dendritic tips of RBC1 are doughnut shaped whereas those of RBC2 ended in a simple tip within the rod spherule, consistent with our light microscopy data (Fig. 3c,d, 5a).

We then focused on one RBC1 and one RBC2 in the central area of the volume and traced all of their post-synaptic neuronal processes (Fig. S3c,d). Amacrine cells (ACs), unlike reginal ganglion cells (RGCs), make output synapses within the retina. Hence, we identified the neuronal class (e.g. AC) or RGC) for the majority (28/32) of the RBC1 postsynaptic processes and over the half (18/31) of the RBC2 postsynaptic processes based on the presence or absence of presynaptic structures. We found that both RBC1 and RBC2 predominantly synapse onto ACs (Fig. 6a,f). The majority of the postsynaptic processes received 4 or fewer ribbon synapses from one RBC1 or RBC2, with an exception of one process, which received 14 inputs from one RBC1 (Fig. 6b,g).

**Figure 6.**
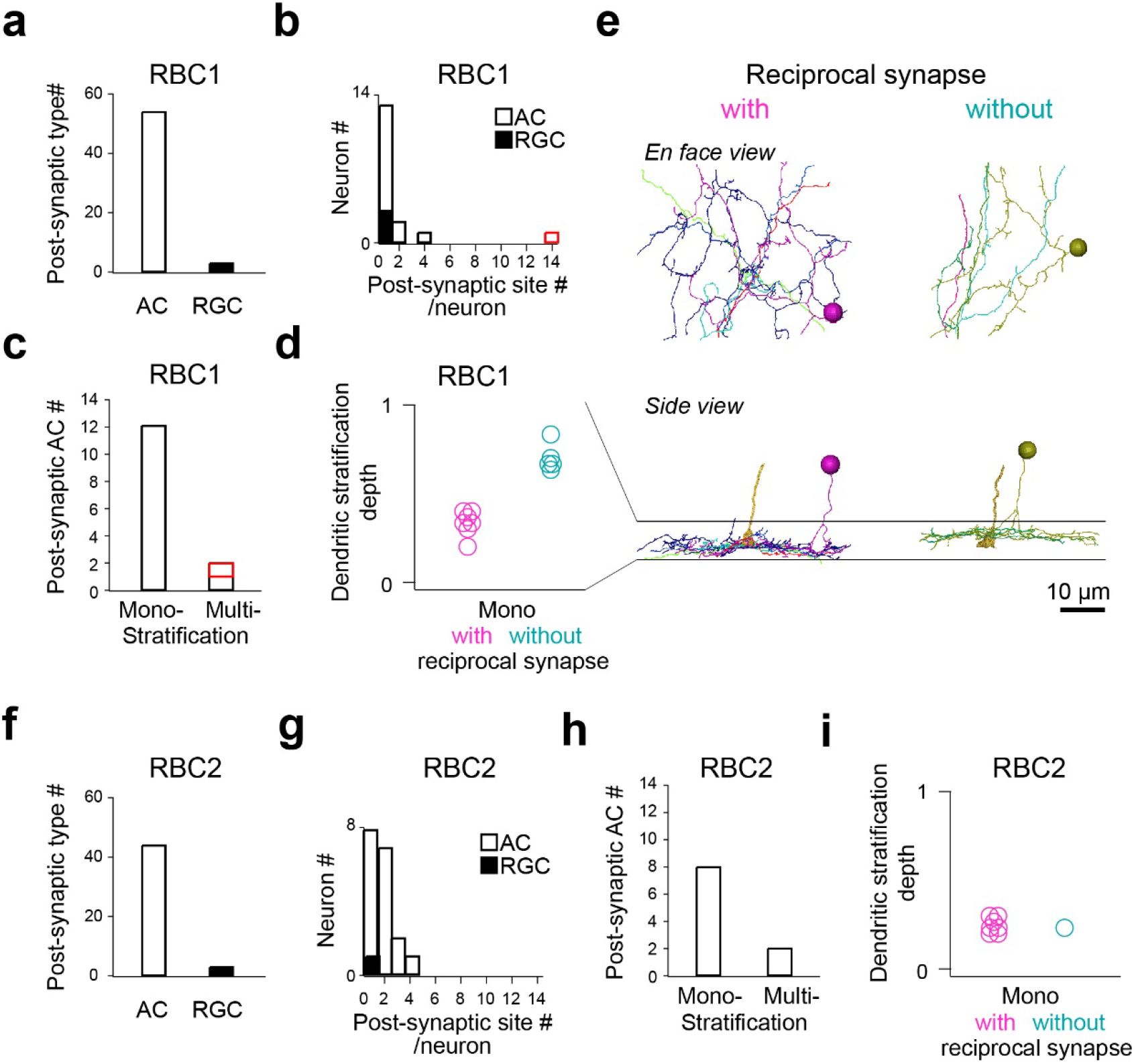
Identification RBC1 and RBC2 post-synaptic neuron types. **a-d**, Quantification of morphological parameters of neurons postsynaptic to one of the RBC1s in the EM volume (marked by an open arrow head in Fig. 5c). One postsynaptic neuron contained an exceptionally higher number of synapses (14) with the RBC1 (marked in red in **b** and **c)**. Dendritic stratification is normalized to 0 and 1 at the lower and upper ends of the RBC1 axon terminals, respectively in **d**. **e,** Mono-stratifying ACs with (red) or without (blue) reciprocal synapses with RBC1s in the volume. The axon of the presynaptic RBC1 is also shown in the side view. Individual cells were color coded. **f-i**, Quantification of morphological parameters for neurons postsynaptic to one of the RBC2s in the EM volume (marked by a closed arrow head in Fig. 5c). AC: amacrine cells, RGC: retinal ganglion cells.

We further morphologically classified the postsynaptic ACs that we traced, comprising 14 cells for RBC1 (Fig. S5-7) and 10 cells for RBC2 (Fig. S8,9), respectively. Most ACs extended their dendrites within a single sublamina in the IPL (Fig. 6c,h). Among these mono-stratifying ACs for RBC1, we identified two groups based on their dendritic stratification depth within the IPL (Fig. 6d,e). These two groups of ACs differed in their synaptic arrangement. ACs stratifying in the lower layer formed reciprocal synapses (RS) – a synaptic arrangement that includes both input and output synapses with a BC axon - with RBC1, whereas ACs stratifying in the upper layer did not (Fig. 6d,e, Fig. S5,6). For RBC2, all but one AC (1/8) formed local reciprocal synapses (Fig. 6i, Fig. S8,9). In addition, RBC1 formed an exceptionally high number of ribbon connections with one bi-stratifying AC (marked in red in Fig. 6b,c, morphology in Fig. 7d,e and Fig. S7). In contrast, RBC2 does not have a post-synaptic partner with extensive synapses.

**Figure 7.**
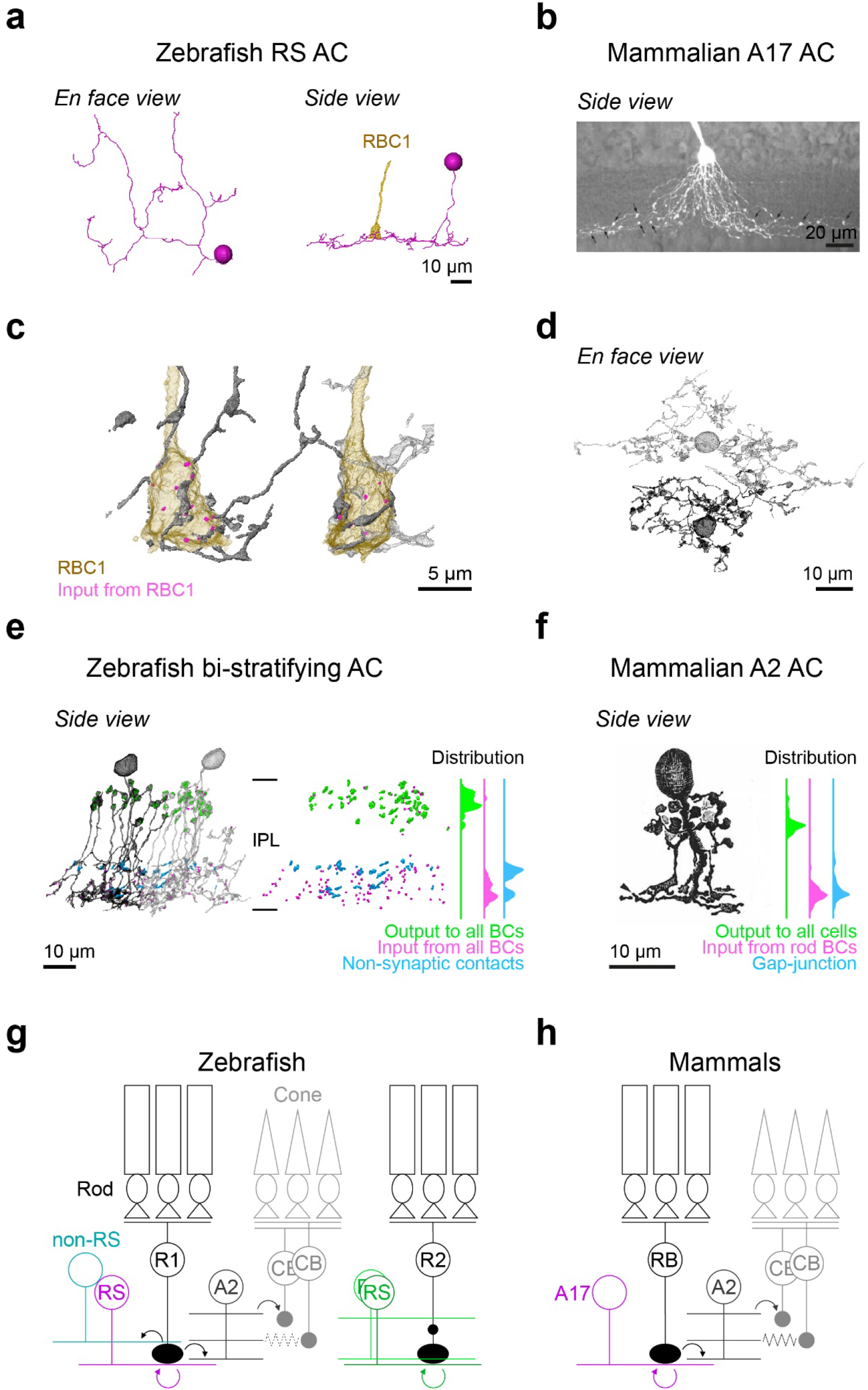
Circuit diagram of RBC1 is similar to mammalian RBC pathway. **a**, An example monostratified amacrine cell (RS AC) classified in Fig. 6e that formed of reciprocal synapse with RBC1. **b,** A mouse A17 amacrine cells (taken from ^47^) (**b**). **c,d,** A2-like ACs that are postsynaptic to two neighboring RBC1s. **e,d**, Locations and distributions of synaptic sites and non-synaptic contacts with BCs in zebrafish bi-stratifying AC (**e**) and in rabbit A2 ACs (taken from ^32^) (**f**). Note that synapses or non-synaptic contacts with AC and RGC are not included in (**e**), and inputs from CBC are not included in (**f**). **g,h,** Schematic diagrams of zebrafish (**g**) and mammalian (**h**) RBC pathways. Mouse A17 and rabbit A2 images are from ^47^ and ^32^, respectively, used with permissions. Data for the distributions of synaptic sites within mammalian A2 ACs across the inner plexiform layer (IPL) are taken from^48^.

Taken together, these results demonstrated that RBC1 allocated the majority (91%) of synaptic outputs to 3 types of AC: 14% to non-RS ACs, 37% to RS ACs, and 40% to one bi-stratifying ACs. In contrast, the RBC2 we reconstructed synapsed primarily onto a mono-stratifying AC type (68%). Among 24 ACs that we traced throughout the volume, only 5 were shared between RBC1 and RBC2 (Fig. S5-9). Therefore, the downstream circuits of these BC types are largely separate at least at the AC level.

### The wiring diagram of RBC1 resembles that of the mammalian primary RBC pathway

Finally, we compared the targets of RBC1 and RBC2 with those known to comprise the primary rod pathway in the mammalian retina. Mammalian RBCs synapse onto a mono-stratifed AC type called A17 and a bi-stratified AC type called A2^3, 5–7^. A17 ACs are wide field ACs that synapse exclusively with RBCs and form reciprocal synapses with them^31, 44^, whereas A2 ACs are narrow field ACs that receive numerous (∼40) synapses from RBCs but do not form reciprocal synapses with them^18, 30^. Instead, A2 ACs form gap-junctions with ON CBCs through dendrites in the ON layer and output synapses onto OFF CBCs through the bouton structures in the OFF layer called lobular appendages^32^.

We found that zebrafish RBC1s synapse onto both wide field ACs with reciprocal synapses (the RS ACs), resembling mammalian A17s, and a narrow field bi-stratifying ACs with extensive synaptic connections, resembling mammalian A2s (Fig. 6b-e, Fig. S5,7). By marking synaptic sites of RS ACs throughout their dendrites, we found that RS ACs are dedicated to the RBC1 pathway; synapsing predominantly (both input and output) with RBC1 and to a lesser extent with RBC2 (Fig. S5), with no synapses with other BC types (n=7 cells). This synaptic specificity and the reciprocal synapse arrangement in RS ACs mirror those of mammalian A17 ACs^31^ (Fig. 7a,b).

In contrast to the targets of RBC1, RBC2 formed synapses exclusively with wide-field ACs (Fig. S8,9) and lack a synaptic partner with extensive synapses. Thus, RBC2 participates in a circuit that differs from that of mammalian RBCs.

## DISCUSSION

By combining scRNA-seq, electrophysiology, and light and electron microscopy circuit reconstructions, we demonstrated that RBC1 shares many features with the mammalian primary rod pathway (Fig. 7g,h), implying that the conserved rod pathway is evolutionarily ancient.

### The number of BC types

In this study, we found 23 molecular types in adult zebrafish BCs. However, a previous morphological characterization of zebrafish BCs, based on their dendritic connectivity with photoreceptors and axon stratifications, identified 32 anatomical types^48^. The discrepancy in the number of BC types between morphological and transcriptional characterizations may arise from the regional specializations that have been documented in the larval zebrafish retina^49–52^. In any event, it is clear that the adult zebrafish retina contains at least 23 BC types.

This number of molecular types of bipolar cells (BCs) in zebrafish, as identified in this study, is higher than that found in mammals investigated to date (14-15 across mammals)^26, 27, 29, 53^, but similar to that found in chick retina (22 molecular and 15 morphological BC types)^54, 55^. The higher number of BC types in zebrafish and chicken is not surprising, given that these species have higher numbers of photoreceptor types: 5 in fish and 7 in chicks, compared to >=3 in mammals^56, 57^. We demonstrate here one source of the increase: a single type of BC carries most of the input from rods in mammals, whereas zebrafish has two RBC types.

### The number of RBC types

Previous morphological characterization of zebrafish BCs found only one BC type that connects rods and red cones. Axons of these BCs terminate in the innermost layer of the IPL^48^. We speculate that this type actually includes both RBC1 and RBC2, which were combined owing to their striking morphological similarity (Fig. 2,3). Consistent with this hypothesis, studies in goldfish have reported two morphologically distinct “mixed” BC types that receive dominant inputs from rods^22^. They have large axon terminals at the bottom of the IPL, but the axon of one mixed BC type contains a smaller axonal distal bouton in the OFF layer, similar to RBC2^58^. Immunostaining for PKC only labels mixed BCs without an axonal distal bouton, similar to RBC1^59^. The presence of these features in goldfish suggests that RBC1 and RBC2 are conserved among teleost fish.

A2- and A17-like ACs may also be conserved in goldfish. Paired elecrophysiological recordings between goldfish RBC1 and ACs revealed that RBC1 provides synaptic inputs to two morphological types of ACs: wide-field mono-stratifying and narrow field bi-stratifying AC types^60^. The dendrites of the bi-stratifying ACs wrap around the RBC1 axon terminals^60^, similar to A2 ACs in zebrafish and mammals (Fig. 7c). Goldfish RBC1 receives GABAergic reciprocal feedback at the axon terminals, like mammalian RBC^61, 62^. Taken together, although it remains unknown whether these two AC types in goldfish exhibit similar synaptic connectivity patterns to those of mammalian A2 and A17 ACs, the findings in goldfish are consistent with the idea that the primary rod pathway, including A2 and A17 ACs, is conserved in goldfish. Some differences in physiological properties between mammalian RBC and goldfish RBC1 were also found. First, goldfish RBC1 receives GABAergic lateral inhibition^61^. The exact cell types that provide this inhibitions are unclear, but it is likely coming from the wide-field mono-stratifying ACs, as their dendrites extend laterally. In contrast, mammalian A17 ACs do not provide lateral inhibition onto RBCs, as each varicosity of A17 ACs at the RBC axon terminals operates independently of each other^63^. Second, goldfish RBC1 exhibits spikes^64, 65^, whereas the spikes are only found in cone BCs in mammals. Nonetheless, the absolute visual sensitivity of goldfish is comparable to that of mammals^66^, suggesting that the primary rod pathway we discovered in teleosts is capable of transmitting information evoked by a single photon.

Much less is known about RBCs in other non-mammalian vertebrate species. In salamander, one type of mixed ON BCs exhibit sensitivity close to that of rods^14, 67^. They terminate their axons at the bottom of the IPL, similar to teleosts and mammals, but it is unknown whether they express the RBC marker PKC. Furthermore, the anatomical connections of BCs with rods are not yet comprehensively studied in salamander. In birds, PKC labels some ON layer stratifying BC types strongly^68, 69^. Single-cell RNA-sequencing of chick BCs revealed a BC type that is transcriptomically similar to the mammalian RBCs^55^. However, the physiological properties and connections of these BCs are unknown. Moreover, a connectomic survey of chick BC types failed to identify BC cells that connect with all rods in their dendritic field^54^. Unlike the species mentioned above, the presumed rod bipolar cells, which have light sensitivity close to that of rods, in sea lamprey are OFF type^70^. However, their connectivity with rods is unknown. Thus, it remains unclear whether RBC2 orthologs are present in species other than zebrafish.

In mammals, morphological, molecular, and functional studies have identified only a single RBC type^23, 26, 27, 71^. Therefore, we speculate that either RBC2 evolved after the divergence between teleost fish and mammals, or mammals lost this pathway.

### Roles of cone inputs in RBCs

Cone inputs onto rod-dominant mixed BCs have been proposed to broaden the dynamic range of light intensities to which they can respond^72^. Consistent with this idea, we found that both RBC1 and RBC2 are selective for red cones, which, with their broad spectral sensitivity, are suited for encoding achromatic luminance information^73^. Because rods evolved from cones^1^, we speculate that RBCs may have emerged from red-cone specific CBCs. The red cone selectivity is also conserved in at least one of the mixed rod dominant BC types in goldfish^74^. Although cone selectivity is unknown in Salamander rod-driven mixed BCs, their spectral sensitivity curve is broader at longer wavelengths than that of rods, indicating that they may connect to red cones^14^.

Electrophysiological recording from rod-driven BCs in *Giant Danio*, a teleost fish species, showed that rod and cone inputs onto rod-dominant BCs are mediated by different mechanisms: rod inputs through mGluR6, whereas cone inputs through both mGluR6 and EAAT (excitatory amino acid transporter)^72^. In this BC type, mGluR6 and EAAT-mediated inputs suppress each other, likely to allow this cell to respond to both rod and cone dynamic ranges^72^. Electrophysiological recordings in zebrafish found that some of ON BCs responded to glutamate via both mGluR6 and a glutamate-gated chloride conductance increase mechanism, which is likely through EAATs^75^. However, the nature of EAAT contributions for cone responses in RBC1 and RBC2 is unknown.

While the study in *Giant Danio* suggest that mixed inputs expand the dynamic range of rod-dominant BCs, electrophysiological recordings in goldfish and salamander have found that the dynamic range of rod dominant BCs is similar to that of rods and that cone contributions to the light response are small^14, 76^. Therefore, the roles of red-cone inputs to RBCs remain to be determined.

### Unifying mixed BCs and RBCs

In mammals, it was initially thought that RBCs exclusively synapse with rods^77^. However, several recent studies have demonstrated convincingly that RBCs also receive synapses from cones, at least in mice and rabbits^8, 77, 78^. Indeed, mouse RBCs contact the majority of M-cones (∼80%), which are analog of zebrafish red-cone, in their dendritic territories^8^. RBCs were likely thought to be exclusive to rods because of the high ratio of rods to cones in the outer nuclear layer in mice and rabbits^79, 80^. As a consequence, only a few cones, generally three or fewer, synapse on a mouse RBC, compared to inputs from ∼35 rods^8, 78^.

The rod-driven BCs in non-mammals are classically called “mixed” BCs because they connect with both rods and cones. However, as argued above, this mixed connectivity is conserved in the mammalian RBCs. Moreover, the dendritic specificity and connectivity of RBCs are conserved in mice and zebrafish. In both species, RBCs connect with all rods and the majority of red-cones (or M-cones) in their dendritic fields (Fig. 3f,h). Therefore, although the coverage of cones in mice is still lower than that in zebrafish, converging rod and red-cone inputs is likely a conserved feature of RBCs in all vertebrates even if the proportion and number of cone inputs may vary across species. This leads us to propose that the non-mammalian mixed BCs and the mammalian RBCs represent a single class of neurons, RBCs. Finally, taken together with the striking similarity in the downstream circuitry of RBCs between zebrafish and mammals, we conclude that zebrafish RBC1 is transcriptomically, anatomically, and functionally equivalent of mammalian RBC and that they share the same evolutionary origin.

## ACKNOWLEDGEMENTS

We thank the Vision Core at the University of Washington for processing zebrafish retina samples and acquiring serial images for SBF-SEM. We thank Rachael N. Swanstrom for helping with cell tracing of the EM volume. Funding was provided by the MCSA fellowship (“ColourFish” 748716) from the EU Horizon 2020 to TY, the NIH EY14358 to ROW, U01MH105960 and R01 EY022073 to JRS, EY01730 (Vision Core grant, P.I. M. Neitz) Developmental Biology Training Grant HD07183 to FDD, The Wellcome Trust 220277/Z20/Z to TB, European Research Council ERC-StG 677687 to TB, UKRI BBSRC, BB/R014817/1 and BB/W013509/1 to TB, Leverhulme Trust PLP-2017-005; RPG-2021-026 to TB, NIH grant R00EY028625 to KS, Hellmann Foundation Fellowship to KS, travel grant from Graduate School in Systemic Neuroscience to YK, and Max Planck Society to YK and HB.

## AUTHOR CONTRIBUTIONS

TY, ROW, AMH, PM, JH, and YK designed the study, with input from KS, JRS, HB, and TB. YK performed single-cell RNA-seq, under the supervision of HB and JRS with guidance from KS. JH processed and analyzed the data with guidance from KS; TY and SCS generated new plasmids; TY and FDD generated novel lines; TY performed experiments, collected and analyzed the data for light microscopy with help from OL; PM performed whole-cell patch recordings; FDD prepared the sample for SBF-SEM; AMH, OL, and TY traced the EM images; TY analyzed the EM data; TY wrote the manuscript with inputs from all authors.

## MATERIALS AND METHODS

### Animals

All procedures were performed in accordance with the University of Washington Institutional Animal Care and Use Committee guidelines, the Harvard University/Faculty of Arts & Sciences Standing Committee on the Use of Animals in Research and Teaching (IACUC), and the UK Animals (Scientific Procedures) act 1986 and approved by the animal welfare committee of the University of Sussex. For all experiments, we used adult zebrafish (age 6-18 months) of either sex that were kept at 28°C in a room with a normal 14/10 light cycles.

The following previously published transgenic lines were used: *Tg(vsx1:GFP)^nns^*^5 83^, *Tg(vsx1:memCerulean)^q^*^19 84^, Tg(*trb2:tdtomato)^q^*^22 85^. In the larva *Tg(vsx1:memCerulean)^q^*^19^ labels a subpopulation of OFF layer stratifying BCs^84^. In adults, while OFF stratifying BCs are still weakly labeled, Cerulean is now strongly expressed in RBC1 (Fig. 2a,c). In addition, *Tg(vsx2:memCerulean)^wst^*^01^ line was generated by injecting pBH-vsx2-memCerulean-pA plasmid into single-cell stage eggs. Plasmid was diluted in 1x Danieau’s solution to a concentration of 50 ng/ml. Plasmid solution was loaded into a pulled-glass micropipette, mounted to a micromanipulator (Narishige), and pressure-injected via attachment to a Picospritzer II (Parker). Injections were made at 10 psi for durations from 100 to 200 ms. Injected fish were raised and out-crossed with wild-type fish to screen for founders. Positive progenies were raised to establish transgenic lines.

### BC single cell RNA sequencing

#### BC purification and sequencing

Adult zebrafish carrying the *Tg(vsx1:GFP)nns5* transgene were used to isolate BCs for single-cell RNA sequencing. Retinas were dissected and digested in papain solution containing 20U/ml papain, 80U/ml DNaseI, and 1.5mM L-cysteine in oxygenated (ox) Ames solution at 28°C for 45 minutes. The digestion was stopped by replacing the papain solution with a papain inhibitor solution containing 15mg/ml ovomucoid and 15mg/ml BSA. The tissue was gently dissociated by trituration using a flamed glass pipette and washed with ox. Ames containing 0.4% BSA. The resulting cell suspension was filtered through a 30μm strainer and fluorescence-activated cell sorting (FACS) was performed. Non-transgenic wild-type retinas were used to determine background fluorescence levels and adjust sorting gates. Live bipolar cells were distinguished using Calcein blue. Cells were washed, resuspended in PBS 0.04% BSA, and loaded onto the microfluidic device within ∼45 minutes after FACS enrichment. Droplet RNA sequencing experiments were conducted on the 10X chromium platform according to the manufacturer’s instructions with no modifications. Up to sixteen retinas from up to eight fish per batch were dissected and dissociated. Eight cDNA libraries were generated across four experiments with two replicates each. The cDNA libraries were sequenced on an Illumina HiSeq 2500 to a depth of ∼30,000 reads per cell.

#### Single cell transcriptomics data analysis

We performed the initial preprocessing using the cellranger software suite (version 2.1.0, 10X Genomics), following steps described previously in our study of Zebrafish RGCs^86^. The sequencing reads were demultiplexed using “cellranger mkfastq” to obtain a separate set of fastq.gz files for each of 8 samples, which were distributed across Y biological replicates. Reads for each sample were aligned to the zebrafish reference transcriptome (ENSEMBL zv10, release 82) using “cellranger count” with default parameters to obtain a binary alignment file and a filtered gene expression matrix (GEM) for each sample. To account for intronic reads, the binary alignment files were processed using velocyto with default parameters^87^, producing a loom file containing a GEM for exonic reads and a separate matrix for intronic reads. The matrices were combined for each sample, resulting in a total gene expression matrix (GEM; genes x cells) summarizing transcript counts. We used the Seurat R package (Stuart et al., 2019) to combine the GEMs from different channels and analyzed them for each of the biological replicates, unless otherwise stated, with default parameter values. Additionally, we evaluated the robustness of our clustering results to variations in select parameters. The full details of our analyses are documented in markdown scripts, which are available at https://github.com/shekharlab/ZebrafishBC.

#### Preprocessing and batch integration

The combined GEM was filtered to remove genes expressed in fewer than 25 cells, and cells expressing fewer than 50 genes resulting in 25,233 genes and 19,492 cells. Briefly, each cell was normalized to a total library size of 10,000 and the normalized counts were log-transformed using the function Seurat::NormalizeData. We used Seurat::FindVariableFetures with option selection.method = “vst” to identify the top 2000 highly variable genes (HVGs)^88^ in each batch. Next, we performed scRNA-seq integration. We used Seurat::FindIntegrationAnchors and Seurat::IntegrateData, both with options “dims=1:40” to perform Canonical Correlation Analysis (CCA)-based batch correction on the reduced expression matrix consisting of the HVGs. The “integrated” expression values were combined across replicates, and used for dimensionality reduction and clustering.

#### Dimensionality Reduction, Clustering and Visualization

To remove scale disparities between genes arising from differences in average expression levels, the integrated expression values for each HVG were z-scored across the cells using Seurat::ScaleData. Next, we performed Principal Component Analysis (PCA) on the scaled matrix, and used Seurat::ElbowPlot to select principal components (PCs). Using the top 20 PCs, we built a k-nearest neighbor graph using Seurat::FindNeighbors and identified transcriptionally distinct clusters using Seurat::FindClusters, using a resolution parameter of 0.5.

Using the top 20 PCs, we also embedded the cells onto a 2D embedding using Uniform Manifold Approximation (Becht et al., 2019) using the Seurat function RunUMAP.

#### Identification of BCs and filtering contaminant classes

BC clusters were identified based on expression of the pan-BC markers *vsx1*, and other cell classes were filtered based on well known gene markers Examples of such genes include *rlbp1a* and *apoeb* for Muller glia^89^, *rbpms2b* for retinal ganglion cells^90^, *gad1* and *gad2* for amacrine cells^91^, *pde6* for photoreceptors^92^, and *cldn19* for endothelial cells^93^. A total of 155 cells corresponding to these cell classes were removed.

#### Hierarchical clustering

To identify transcriptional relationships between BC clusters, we used Seurat::FindVariableFeatures to recalculate the top 2000 most variable genes. The average expression values of genes in each cluster were used as input for hierarchical clustering, performed using pvclust with parameters method.hclust = "complete" and method.dist = "correlation". The resulting output was visualized as a dendrogram.

### Plasmid construction

Plasmid pBH-vsx2-memCerulean-pA was made using the Gateway system (ThermoFisher, 12538120) with combinations of entry and destination plasmids as follows: pTol2CG2 ^94^, p5E-vsx2, pME-membrane-Cerulean, p3E-pA (Kwan et al., 2007). Plasmid p5E-vsx2 was generated by inserting a polymerase chain reaction (PCR)-amplified vsx2 promoter genomic fragment into p5E plasmid using BP clonase (ThermoFisher, 11789013). PCR reaction was performed using primers: 5’-GGGGACAACTTTGTATAGAAAAGTTGATGCTAAACAACTTCAAACGACCAA-3’ and 5’-GGGGACTGCTTTTTTGTACAAACTTGGCCTCTGAGACTATTCCCTTCTTTG-3’.

### Immunostaining and light microscopy imaging

Adult zebrafish were humanely euthanized in ice-chilled fish water. After decapitation, retinal tissues were dissected from the enucleated whole eyes by removing cornea, lens, and epithelial layer in 1x in phosphate-buffered saline (PBS). The tissue were immediately fixed in 4% paraformaldehyde (Agar Scientific, AGR1026) in PBS for 20 min at room temperature (RT) followed by three washes in PBS. For retinal slice preparation, the tissues were mounted in 2% agarose in PBS and sliced at 100 or 200 µm thickness using vibratome (TPI 1000). For rod staining, the tissues were sliced horizontally, parallel to the outer plexiform layer (OPL), to facilitate antibody penetration in the tissue while preserving bipolar cell dendrites in the OPL. Sliced or the whole retinal samples were treated with PBS containing 0.2% Triton X-100 (Sigma-Aldrich, X100) for at least 10 min and up to 1 day, followed by the addition of primary antibodies. After 3 to 5 days of incubation at 4°C, samples were washed three times with PBS and 0.2% Triton X-100 solution and treated with secondary antibodies. After 1 day of incubation, samples were mounted in 1% agar in PBS on a coverslip, and subsequently, PBS was replaced with mounting media (VECTASHIELD, H-1000) for imaging.

Primary antibodies were 4C12 antibody (mouse, 1:50; kindly provided by Jim Fadool ^95^ and anti-mGluR6b antidoby (rabbit, 1:500; kindly provided by Stephan CF Neuhauss ^96^). Secondary antibodies were AlexaFluor594 anti-rabbit (donkey, 1:500; Jackson ImmunoResearch Laboratories 711-586-152) and DyLight647 anti-mouse (donkey, 1:500; Jackson ImmunoResearch Laboratories 715-606-150). Confocal image stacks were taken on a TCS SP8 (Leica) with a 63× oil immersion objective (HC PL APO CS2, Leica). Typical voxel size was 90 nm and 0.5 μm in xy and z, respectively. Super-resolution images were taken on an OMX (General Electric). Contrast, brightness, and pseudo-color were adjusted for display in Fiji [National Institutes of Health (NIH)].

Image stacks were median filtered in Fiji. For some images, maximum-intensity projections were generated in Amira (FEI). 3D image reconstructions were digitally sliced using the Amira slice functions. All measurements were made in Fiji.

### Image Analysis

#### Quantitation of cell density

We obtained the cell density of RBC1 and RBC2 by counting the axon terminals of these cells within regions of interest from confocal image stacks of the dorsal and ventro-temporal retina. RBC2 was counted from images of the *Tg(vsx2:memCerulean)^wst^*^01^ line. Because not all RBC1 labeled in the *Tg(vsx1:memCerulean)^q^*^19^ line express mCerulean, we quantified the density of PKC labeled cells with axon terminals in the bottom layer of the IPL. Counts were obtained from 3-4 retinas from 3 animals of each line. For RBC1, axons were quantified within an area between 37,000 and 85,000 µm^2^, and for RBC2, the areas were between 11,000 -22,000 µm^2^.

#### Dendritic field

The dendritic field was defined by tracing the extent of a given cell’s dendrites with the polygonal select tool, and removing any concavity using FIJI (see Figure S2). The dendritic arbor area was then obtained by calculating the area enclosed by the polygon. Because the dendritic tips of some neighboring cells of the same type overlapped and could not be distinguished readily, one investigator repeatedly traced (3 to 4 measurements for a single cell) the dendrite boundary, and obtained the respective area for a given cell until at least three measurements were within ± 2.5% of the average of all previous measurements for that cell. Confocal images from three fish were used, with images of RBC1 and RBC2 cells acquired from the dorsal and ventral regions of the retina: 10 to 17 cells per fish were measured for each location and cell type, resulting in a total of 33 to 41 cells measured for each location and cell type.

#### Photoreceptor connectivity

Dendritic contacts with photoreceptors were defined by the co-localization of dendritic tips extending towards outer nucleus layer and photoreceptor terminals by scrolling through confocal image stack in Fiji. The percentage contacted was computed by dividing the number of photoreceptors contacted by a given BC by the number of photoreceptors within the dendritic field of the BC.

### Electrophysiology

Fish (3-6 months old) used in physiology experiments were dark adapted for at least 2 hours, and the retinas were isolated under infrared light following procedures approved by the Administrative Panel on Laboratory Animal Care at the University of Washington. Retinas were continuously superfused (∼8 mL/min) with oxygenated (95% O2, 5% CO2) bicarbonate-buffered Ames solution (Sigma) maintained at 25°C–28°C.

Recordings were conducted in a flat-mount preparation with photoreceptors facing down. Bipolar cells were patched at the axon terminals. To access bipolar cell terminals for recording, small groups of ganglion cells were suctioned off the top of the retina to expose the inner plexiform layer. Terminals of RBC1 bipolar cells could be targeted for recording using only infrared illumination, whereas RBC2 terminals, which were not easily visible without fluorescence imaging, were targeted using a custom-built two-photon microscope. As a result, measuring rod-mediated responses in RBC2 was unfeasible due to the compromise of rod responses by two-photon imaging.

Whole-cell voltage-clamp recordings were obtained using patch pipettes filled with a Cs+-based internal solution. This internal solution also included Alexa Fluor 594, which was used for two-photon imaging of each cell after recording to confirm its type by morphology. To isolate excitatory postsynaptic currents, we voltage-clamped cells near the reversal potential for chloride-mediated conductances (∼-60 mV). In addition, to block inhibitory synaptic transmission, we added the GABAA receptor antagonist gabazine (20 μM), the GABAC receptor antagonist TPMPA (50 μM), and the glycine receptor antagonist strychnine (3 μM) to the superfusion solution. In experiments in which mGluR6-mediated input was blocked, the mGluR6 receptor agonist APB (10 μM) was also added to the superfusion.

Light from blue or red light-emitting diodes (LEDs, peak output = 470 nm and 640 nm respectively) was delivered to the recording chamber via fiber optic cable positioned beneath the microscope’s condenser lens. The light uniformly illuminated a circular area through an aperture 0.5 mm in diameter centered on the recorded cell. Protocols for light stimulation were designed to either activate rods only (using the blue LED) or both rods and cones (using the red LED).

### EM data acquisition, reconstruction, and annotation

Dissected retinal tissues from wild type adult zebrafish were immediately transferred into a 1.5-ml tube with the fixative (4% glutaraldehyde in 0.1M cacodylate buffer [pH7.4]) and incubated overnight on a shaker at RT. Subsequently, the tissue was washed three times in 0.1 M cacodylate buffer (pH7.4) and incubated in a solution containing 1.5% potassium ferrocyanide and 2% osmium tetroxide (OsO4) in 0.1 M cacodylate buffer [0.66% lead in 0.03 M aspartic acid (pH 5.5)] for 1 hour. After washing, the tissue was placed in a freshly made thiocarbohydrazide (TCH) solution (0.1 g of TCH in 10 ml of double-distilled H2O heated to 600°C for 1 hour) for 20 min at RT. After another rinse, at RT, the tissue was incubated in 2% OsO4 for 30 min at RT. The samples were rinsed again and stained en bloc in 1% uranyl acetate overnight at 40°C, washed, and stained with Walton’s lead aspartate for 30 min. After a final wash, the retinal pieces were dehydrated in a graded ice-cold alcohol series and placed in propylene oxide at RT for 10 min. Last, the sample was embedded in Durcupan resin. Semithin sections (0.5 to 1 μm thick) were cut and stained with toluidine blue, until the fiducial marks (box) in the ganglion cell layer (GCL) appeared. The block was then trimmed and mounted in a serial block-face scanning electron microscope (GATAN/Zeiss, 3View). Serial sections were cut at a thickness of 70 nm and imaged at an xy resolution of 7 nm. Six tiles, each about 40 μm by 40 μm with an overlap of about 10%, covering from the outer nucleus layer to the ganglion cell layer in a side view was obtained. Retinal location was not recorded. The image stacks were concatenated and aligned using TrakEM2 (NIH). Neurons were traced or painted using the tracing and painting tools in TrakEM2.

### Statistics

Mann-Whitney U test was used to determine the *p*-value for comparing dendritic field sizes.

## DATA AVAILABILITY

Computational scripts detailing scRNA-seq analysis reported in this paper are available at https://github.com/shekharlab/

ZebrafishBC. We have also provided R markdown (Rmd) files that show step-by-step reproduction of the key results at https://github.com/shekharlab/ZebrafishBC. The raw and processed scRNA-seq data reported in this paper was obtained from the Gene Expression Omnibus (GEO) entry GSE237215 (subseries GSE237214).

---

**Figure S1.**
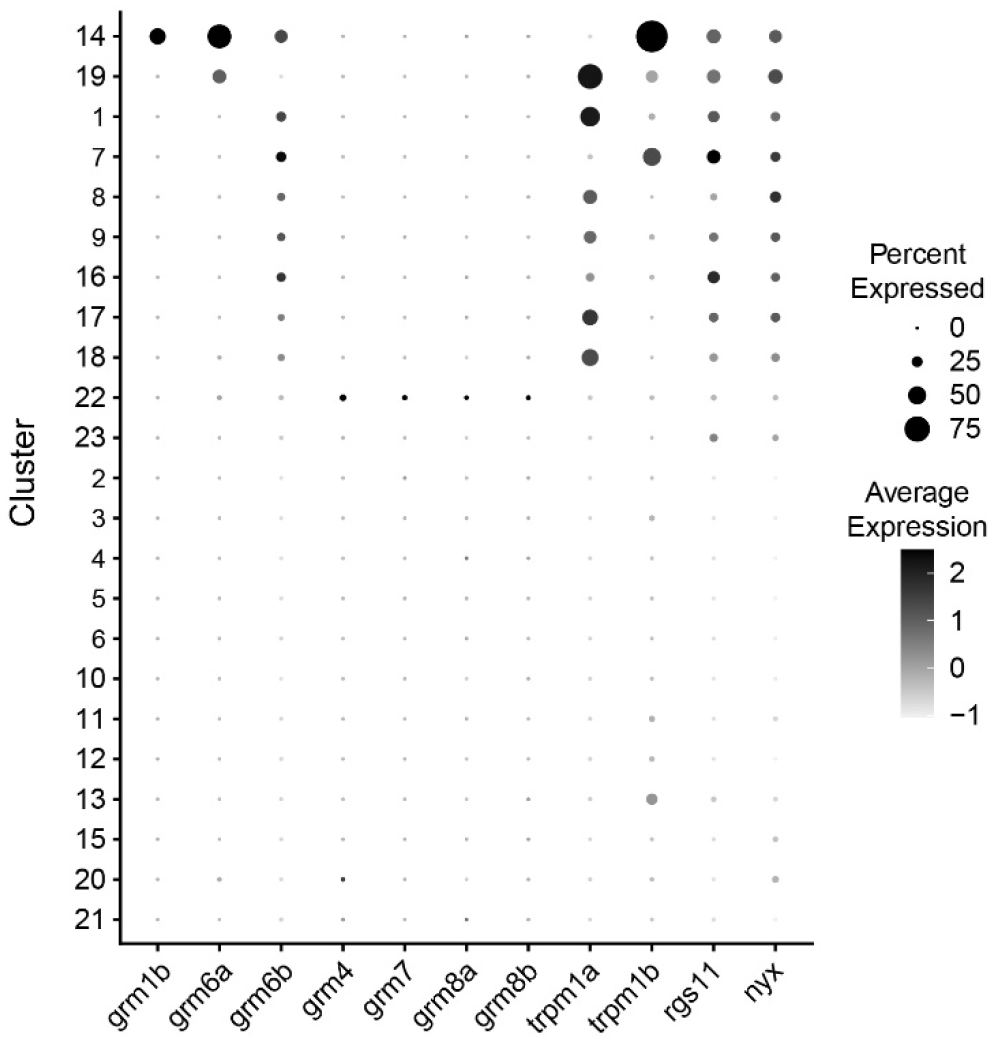
Expression patterns of the identified marker genes in BC clusters. The size of each circle depicts the percentage of cells in the cluster in which the marker was detected (≥1 UMI), and its grey scale depicts the scaled average expression level of cells within the cluster

**Figure S2.**
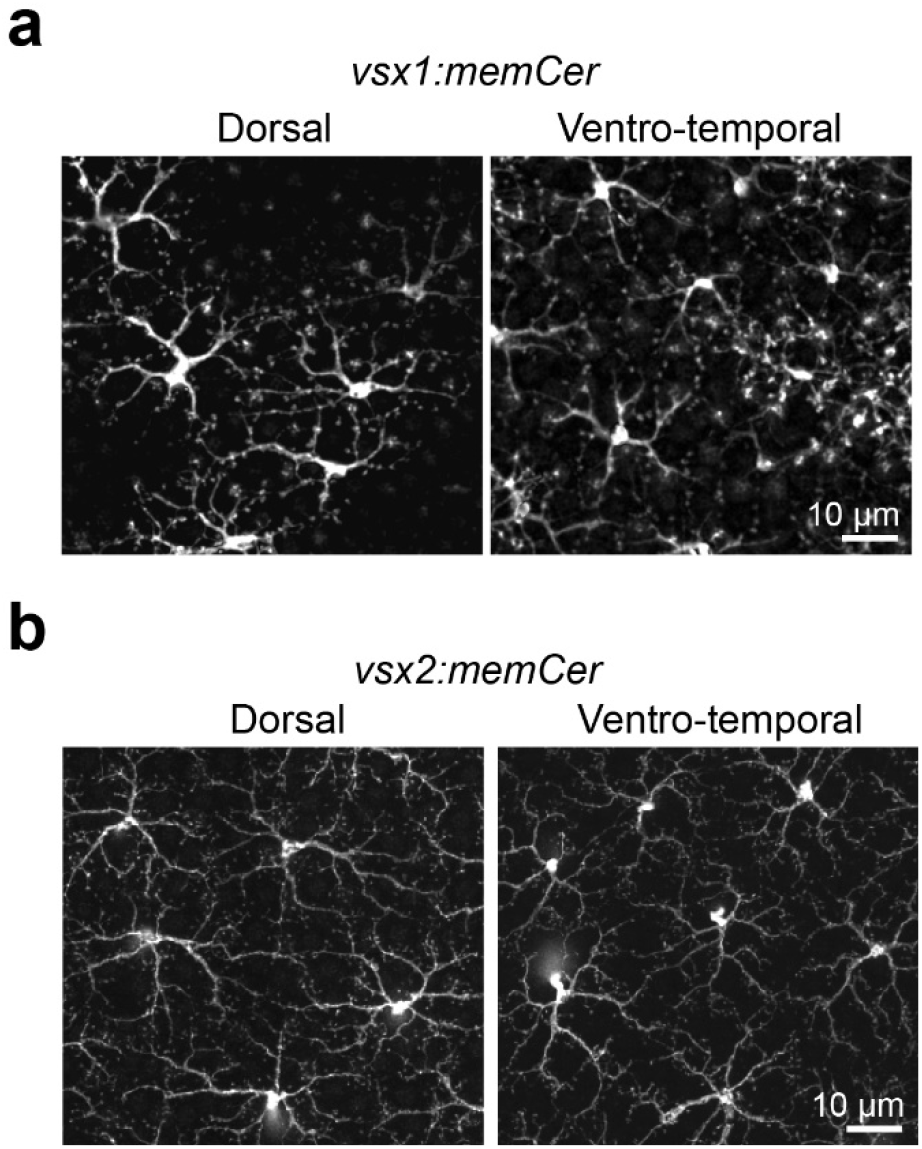
Dendritic tiling of RBC1 and RBC2 BCs across the retina. Confocal images of retinal flat mount at outer plexiform layer level from Tg(vsx1:memCerulean)^q19^ (vsx1:memCer) and Tg(vsx2:memCerulean)^wst01^ (vsx2:memCer). Note that the vsx1:memCer line occasionally labels OFF BCs. These BCs were distinguished by tracing the cells to the axon terminals in the confocal image volumes.

**Figure S3.**
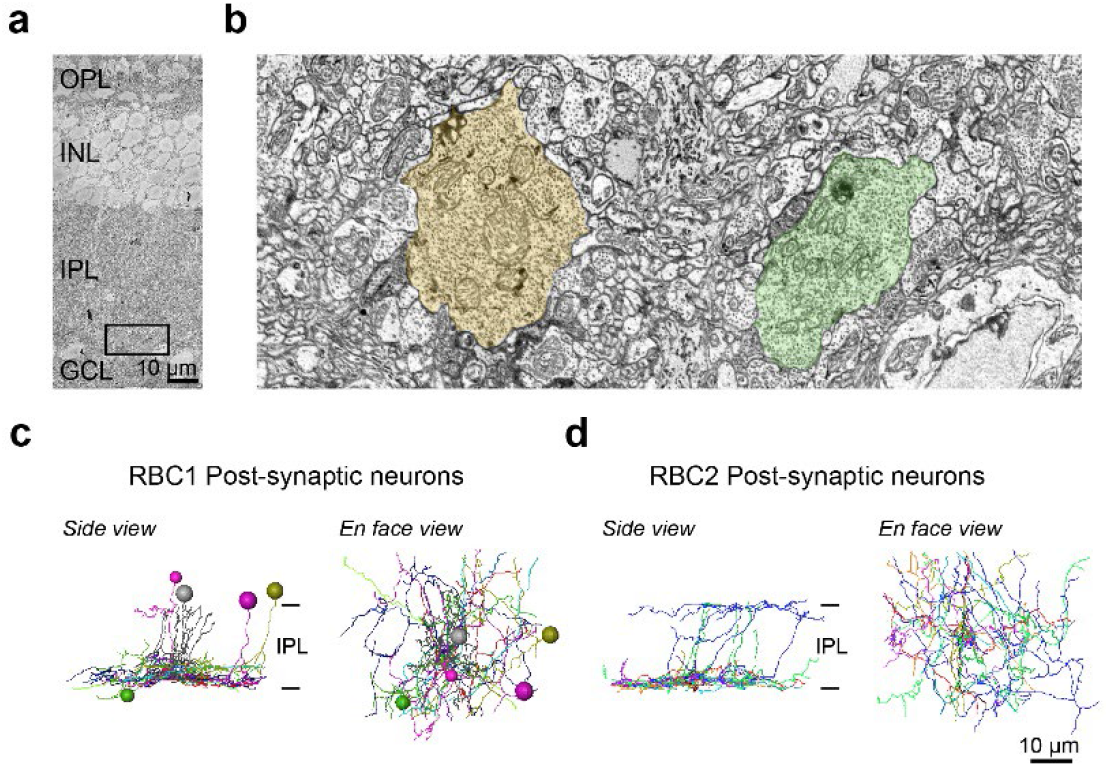
Identification of RBC1 and RBC2 postsynaptic neurons in SBFSEM volume. **a**, A partial image of an example SEM image of an adult zebrafish retina. OPL (outer plexiform layer), INL (inner nuclear layer), IPL (inner plexiform layer), GCL (ganglion cell layer). **b**, Magnified image of the region within the black box in **a** at the bottom layer of the IPL. Characteristic large bipolar cell axons are painted in light yellow and green. **c,d,** Traces of neuronal processes and the location of somas of cells that are post-synaptic to RBC1 and RBC2 cells. Individual cells were color coded. IPL: inner plexiform layer.

**Figure S4.**
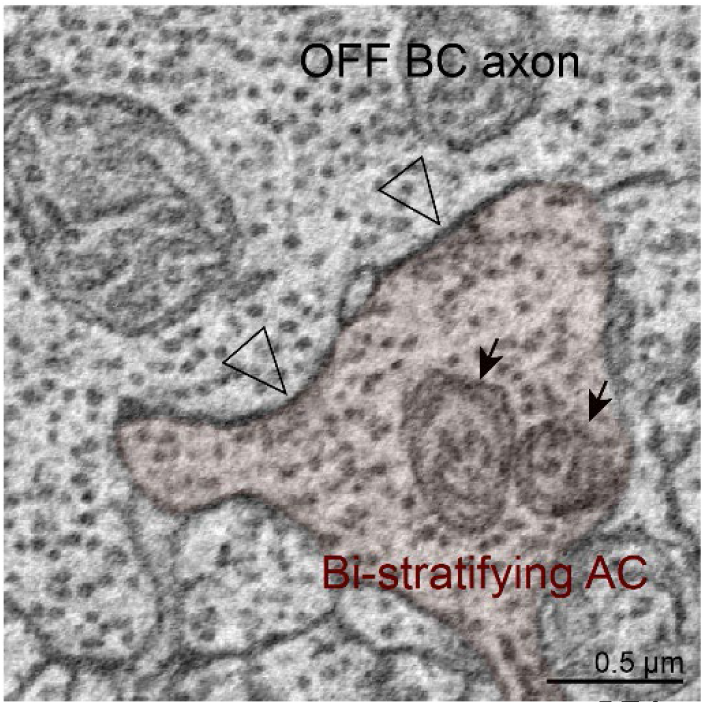
Ultrastructure of the zebrafish A2 AC boutons in the OFF layer. Examples of large synaptic sites (arrowheads) between bi-stratifying A2-like AC and OFF BC axon terminals in the OFF layer. The A2 AC boutons often contain mitochondria (arrows). Next, we examined the synaptic arrangements of RBC1 with the A2-like ACs. First, we confirmed that this type of AC is common to other RBC1s. By tracing the postsynaptic processes of neighboring RBC1, we found another A2-like AC, which received a high number of ribbon inputs from the neighboring RBC1 (Fig. 7c,d). We marked the locations of synapses with all BCs for those two ACs (Fig. 7e). Gap-junctions are too small to be resolved in our SBFSEM images, but as a proxy, we marked non-synaptic contacts (Fig. 7e). These revealed a striking similarity in the distribution patterns of synapses and (potential) gap-junction sites across the IPL layers between this AC type in zebrafish and mammalian A2 ACs (Fig. 7e,f), including the bouton structures in the OFF layer that contain large presynaptic sites and mitchondria^45–47^ (Fig. S4). Taken together, we conclude that the circuit diagram among mammalian RBC, A2, and A17 ACs are conserved in the zebrafish RBC1 pathway (Fig. 7g,h).

**Figure S5.**
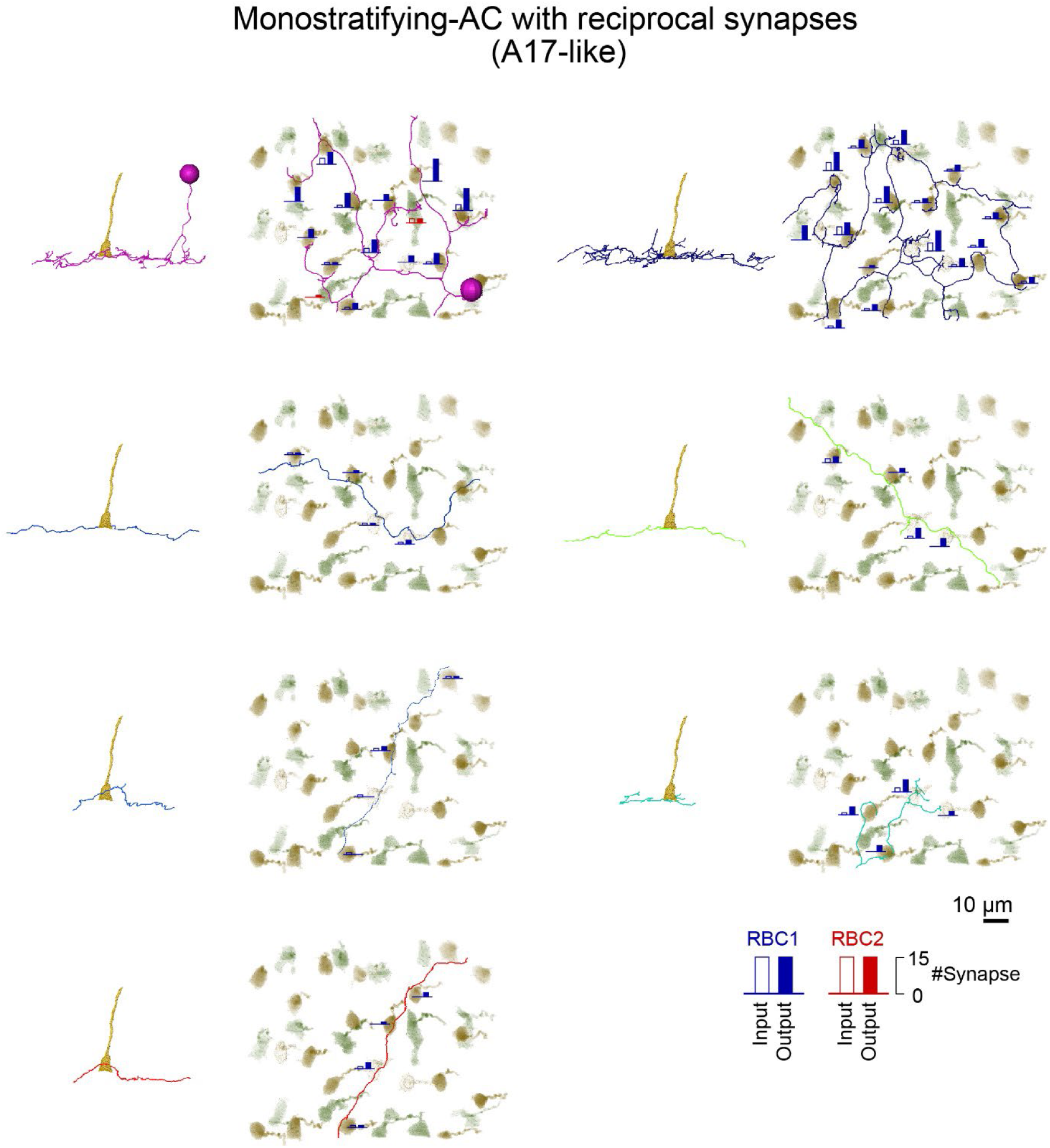
Gallery of mono-stratifying AC making reciprocal synapses with RBC1. *En face* and side views of individual cells or processes. The numbers of input (open bar) and output (closed bar) synapses with each RBC1 (blue) and RBC2 (red) terminal are indicated as the height of the bars.

**Figure S6.**
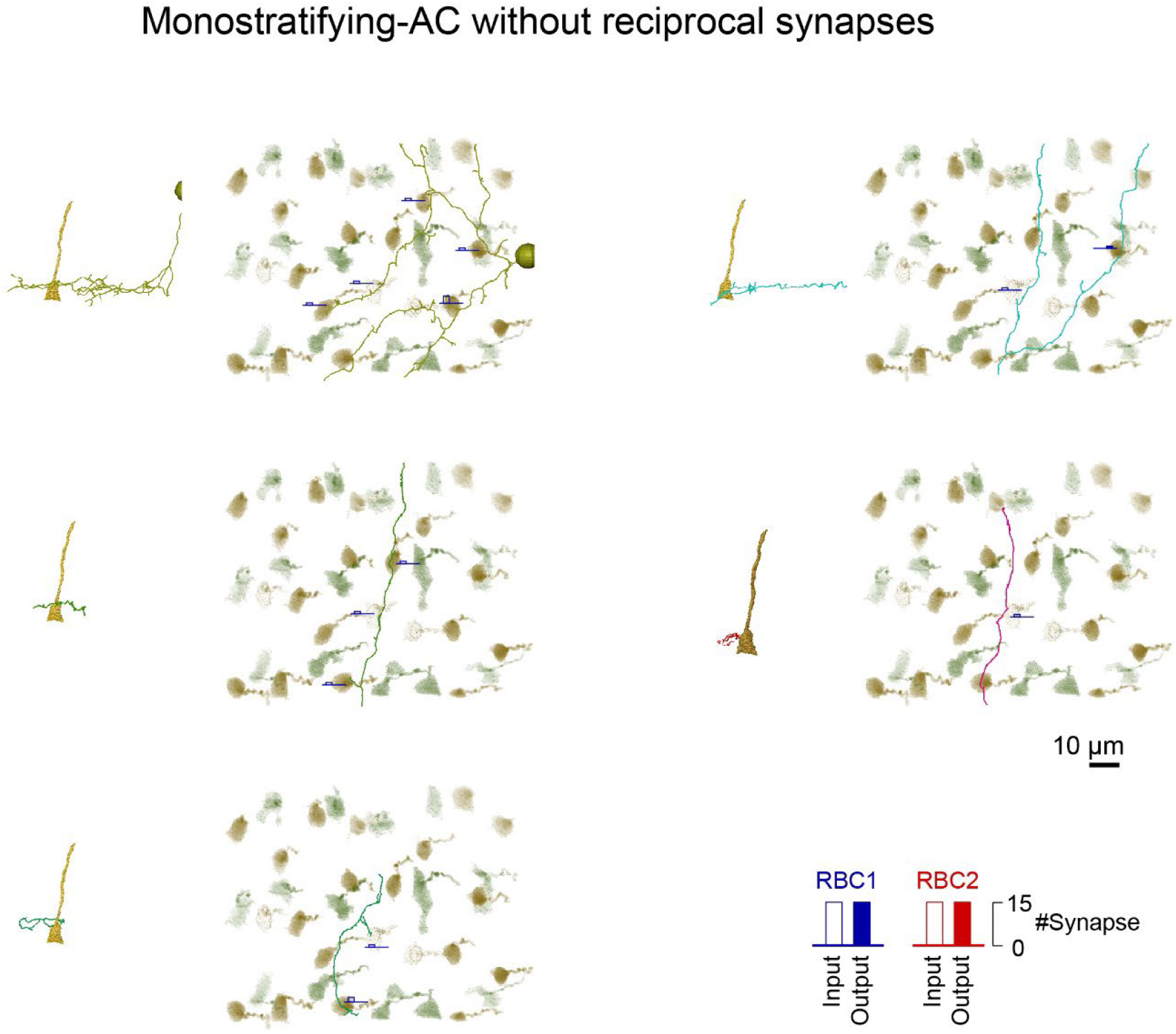
Gallery of mono-stratifying AC without reciprocal synapses with RBC1. *En face* and side views of individual cells. The numbers of input (open bar) and output (closed bar) synapses with each RBC1 (blue) and RBC2 (red) terminal are indicated as the height of the bars.

**Figure S7.**
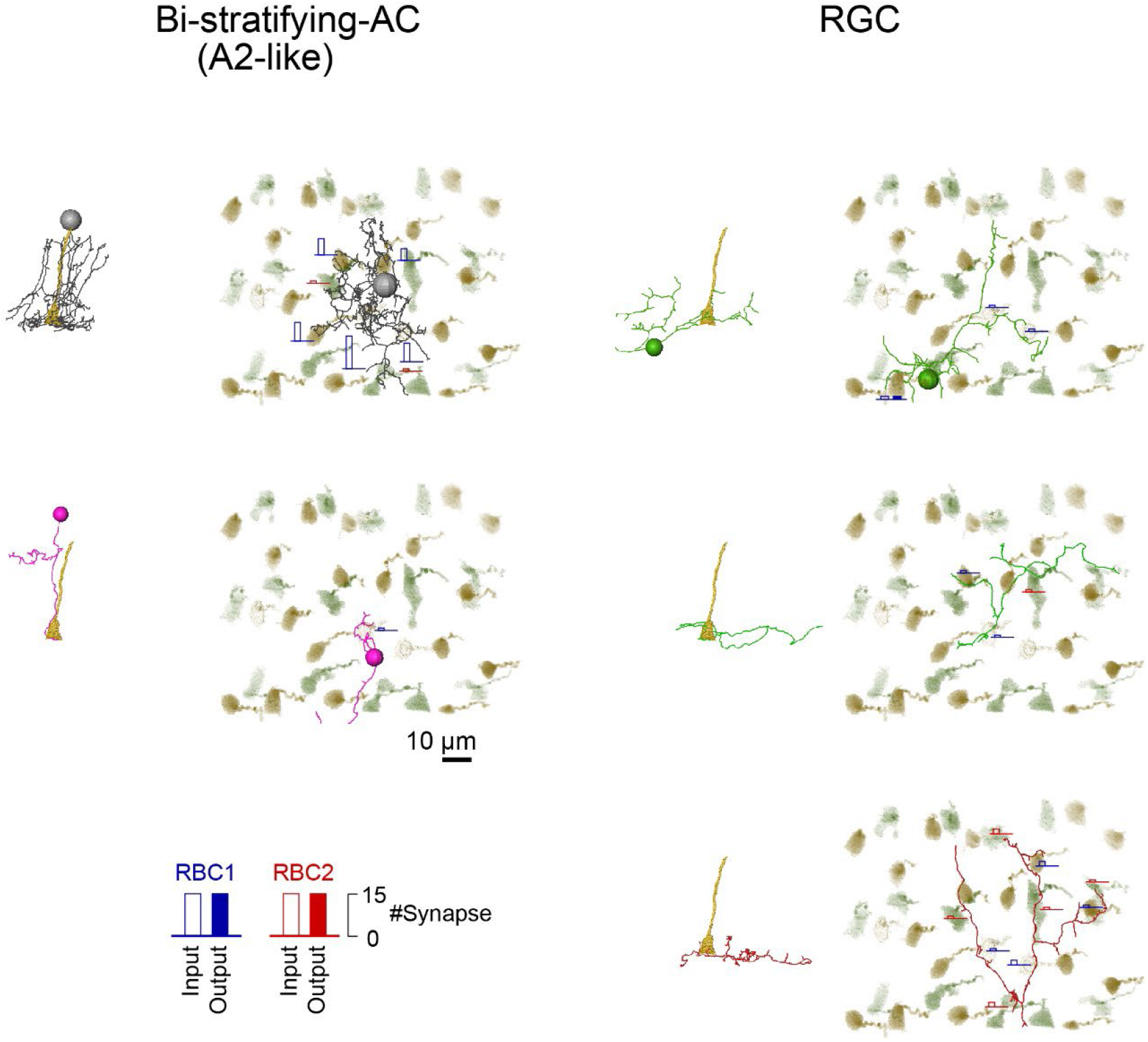
Gallery of bi-stratifying AC and RGC contacted to RBC1. *En face* and side views of individual cells. The numbers of input (open bar) and output (closed bar) synapses with each RBC1 (blue) and RBC2 (red) terminal are indicated as the height of the bars.

**Figure S8.**
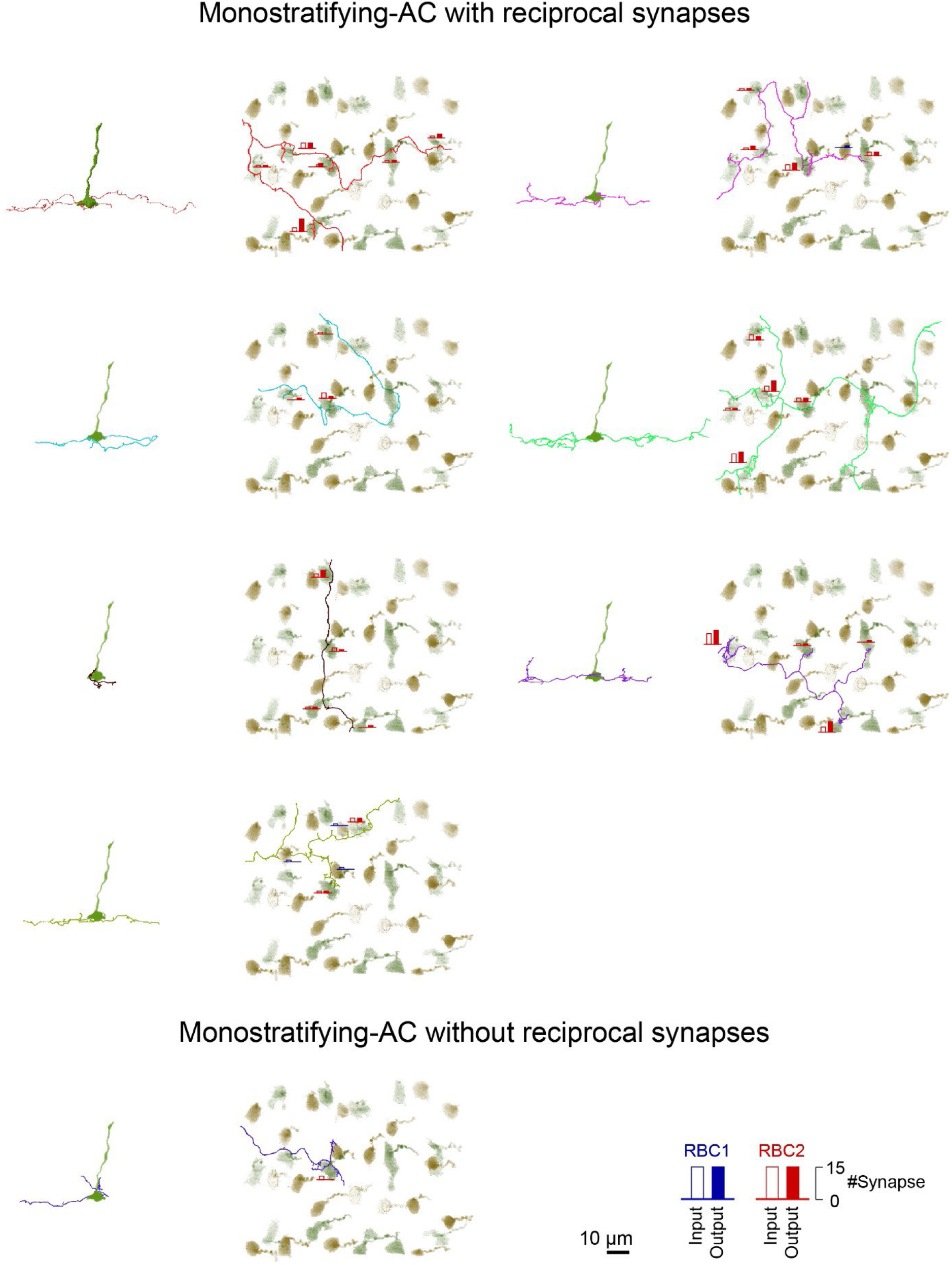
Gallery of mono-stratifying AC connected to RBC2. *En face* and side views of individual cells. The numbers of input (open bar) and output (closed bar) synapses with each RBC1 (blue) and RBC2 (red) terminal are indicated as the height of the bars.

**Figure S9.**
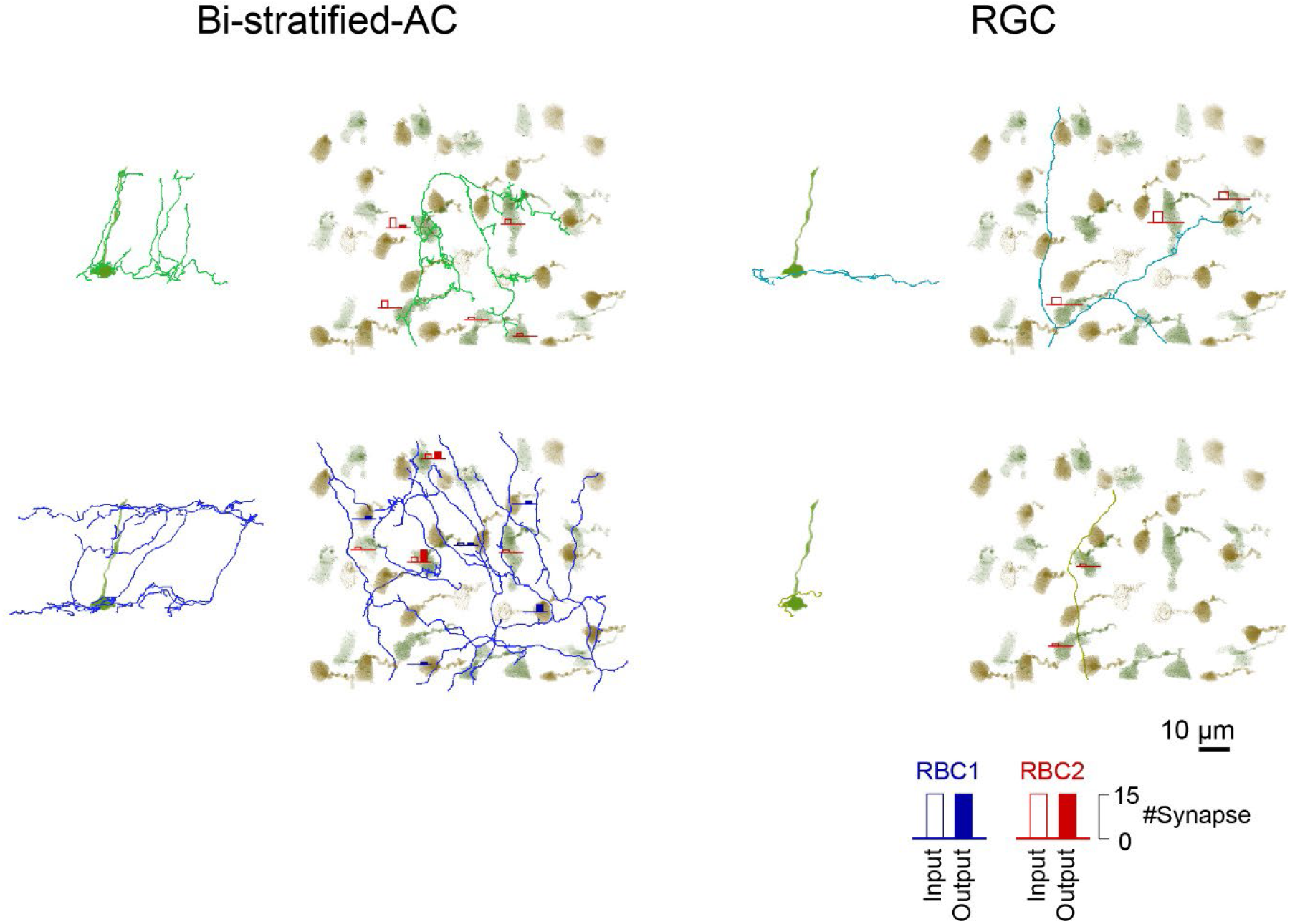
Gallery of bi-stratifying AC and RGC connected to RBC2. *En face* and side views of individual cells. The numbers of input (open bar) and output (closed bar) synapses with each RBC1 (blue) and RBC2 (red) terminal are indicated as the height of the bars.

